# Direct Observation of Competing Prion Protein Fibril Populations with Distinct Structures and Kinetics

**DOI:** 10.1101/2022.08.10.503301

**Authors:** Yuanzi Sun, Kezia Jack, Tiziana Ercolani, Daljit Sangar, Laszlo Hosszu, John Collinge, Jan Bieschke

## Abstract

In prion diseases, fibrillar assemblies of misfolded prion protein (PrP) self-propagate by incorporating PrP monomers. Using total internal reflection and transient amyloid binding super-resolution microscopy, our study analyses elongation of single PrP fibrils to reveal polymorphic populations, featuring structural and dynamic heterogeneity similar to prion strains, which were previously hidden in ensemble measurements. PrP fibrils elongated along a preferred direction by an intermittent ‘stop-and-go’ mechanism. Fibrils fell into three main populations, which each displayed distinct elongation mechanisms incorporating different monomer structures and which maintained their properties even under elongation conditions favouring a different fibril type. Elongation of RML and ME7 prion rods likewise exhibited unique kinetic features. The discovery of polymorphic fibril populations of amyloid and prions growing in competition suggests that prions may present as quasispecies of structural isomorphs and that the replication environment may tilt the balance between prion isomorphs and amyloid species.

**Highlights:**

- Synthetic prion fibril populations contain structurally distinct fibril types
- Fibril types faithfully elongate by different mechanisms
- Fibril types compete for substrate depending on environment
- Fibril populations model quasi-species behavior of prion strains

**eTOC:** Replication of different prion strains causes distinct disease phenotypes. Sun et al. analyzed the growth of individual synthetic prion protein fibrils by super-resolution microscopy and found populations of structurally distinct fibril types, which grew in competition to each other as a quasi-species, recapitulating basic prion strain characteristics *in vitro*.

## Introduction

The abnormal aggregation of cellular proteins into insoluble amyloid is associated with a number of human diseases (Dobson et al., 2020), among them neurodegenerative disorders such as Alzheimer’s Disease (AD), Parkinson’s Disease (PD) and prion diseases (Iadanza et al., 2018). In prion diseases, benign cellular prion protein (PrP^C^) is converted to fibrillar assemblies of misfolded PrP, which self-propagate by recruiting PrP monomers (Collinge, 2016; Prusiner, 1998). This autocatalytic process defines the infectivity of prions (Bieschke et al., 2004). During the misfolding process, PrP converts from α-helices rich folded monomer to amyloid fibrils stacked by parallel in-register β-sheets (Kraus et al., 2021; Manka et al., 2022a; Manka et al., 2022b), but how the structural conversion takes place is not well understood on a molecular level.

In amyloid formation via nucleated polymerization mechanisms (reviewed in (Eichner and Radford, 2011; Iadanza et al., 2018)), fibril elongation is responsible for the increase of fibril mass and is generally the fastest assembly step. Recombinant PrP^C^ readily forms amyloid fibrils under partially denaturing conditions *in vitro*, and the elongation kinetics have been systematically studied by seeding assays in bulk (Honda and Kuwata, 2017; Milto et al., 2014; Ziaunys et al., 2021). Under these conditions, fibrils grow via the addition of unfolded monomers, yielding a mechanism similar to Michaelis–Menten enzyme kinetics (Honda and Kuwata, 2017). Unlike natively unfolded proteins, native folded PrP^C^ can inhibit elongation (Honda and Kuwata, 2017). However, bulk reactions cannot capture the variations in assembly caused by heterogeneous fibril structures, such as seen in prion strains (Collinge and Clarke, 2007).

Aβ fibrils formed *in vitro* (Gremer et al., 2017; Petkova et al., 2005) were shown to adopt different structures depending on incubating conditions. *Ex vivo* Aβ42 fibrils exhibited distinct structures with a direct link to disease type (Yang et al., 2022). PrP fibrils formed *in vitro* showed a high level of polymorphism even under the same incubation condition (Alvarez-Martinez et al., 2011). Structural polymorphism is also observed for authentic prions. Fibrils from different prion strains, which are associated with distinct clinical and neuropathological features (Mead et al., 2014), were shown to have distinct misfolded PrP structures (Collinge, 2016; Kraus et al., 2021; Manka et al., 2022a; Manka et al., 2022b; Safar et al., 1998).

Emerging studies analyse fibril elongation kinetics on a single-particle level in real-time using atomic force or optical microscopy, revealing fine-grained growth properties, such as stop-and-go patterns, growth polarity and fibril breakage (Ban et al., 2003; Ban et al., 2004; Sang et al., 2018; Watanabe-Nakayama et al., 2020; Wordehoff et al., 2015; Young et al., 2017). We adapted total-internal-reflection (TIRF) and transient amyloid binding (TAB) super-resolution microscopy to analyse fibril elongation kinetics of synthetically formed recombinant mouse PrP 91 – 231 (MoPrP 91) fibrils and authentic prion rods on a single-particle level. Our study shows that fibrils grow by an intermittent ‘stop-and-go’ mechanism in a preferred elongation direction. Fibrils formed under seemingly homogeneous conditions can seed multiple fibril types with characteristic structures, which are able to elongate at distinct rates following distinct mechanisms. The fastest-growing fibrils recruited unfolded PrP and displayed a Michaelis-Menten-like mechanism with inhibition at high PrP concentration. The elongation of different types of fibrils was favoured under specific conditions, but fibril types faithfully templated their respective structures even under unfavourable conditions, resembling prion ‘strains’. The elongation kinetics of two prion strains, RML and ME7, *in vitro* likewise exhibited unique kinetic features.

## Results

### Real-time kinetic analysis of single fibril growth

For elongation experiments, fibrillar MoPrP 91 seeds were generated through sequential seeding (Figure S1A, B) and deposited onto the bottom of an inverted microscope chamber. The unbound seed was removed and fibril elongation was monitored by TIRF microscopy over a range of PrP^C^ concentrations (0.3 – 10 μM), GdnHCl concentrations (1.4 – 2.3 M) and temperatures (27 – 40°C) in the presence of Nile Blue (400 nM) as the amyloid-specific probe. Dot-like seeds grew gradually into long single fibrils (Figure 1A; a) or multiple fibrils in different directions (b), indicating that those dots were seed clusters. To analyze the kinetics of elongation, a kymograph was generated for each fibril (Figure 1B) and converted to a fibril growth plot (Figure 1C).

**Figure 1.**
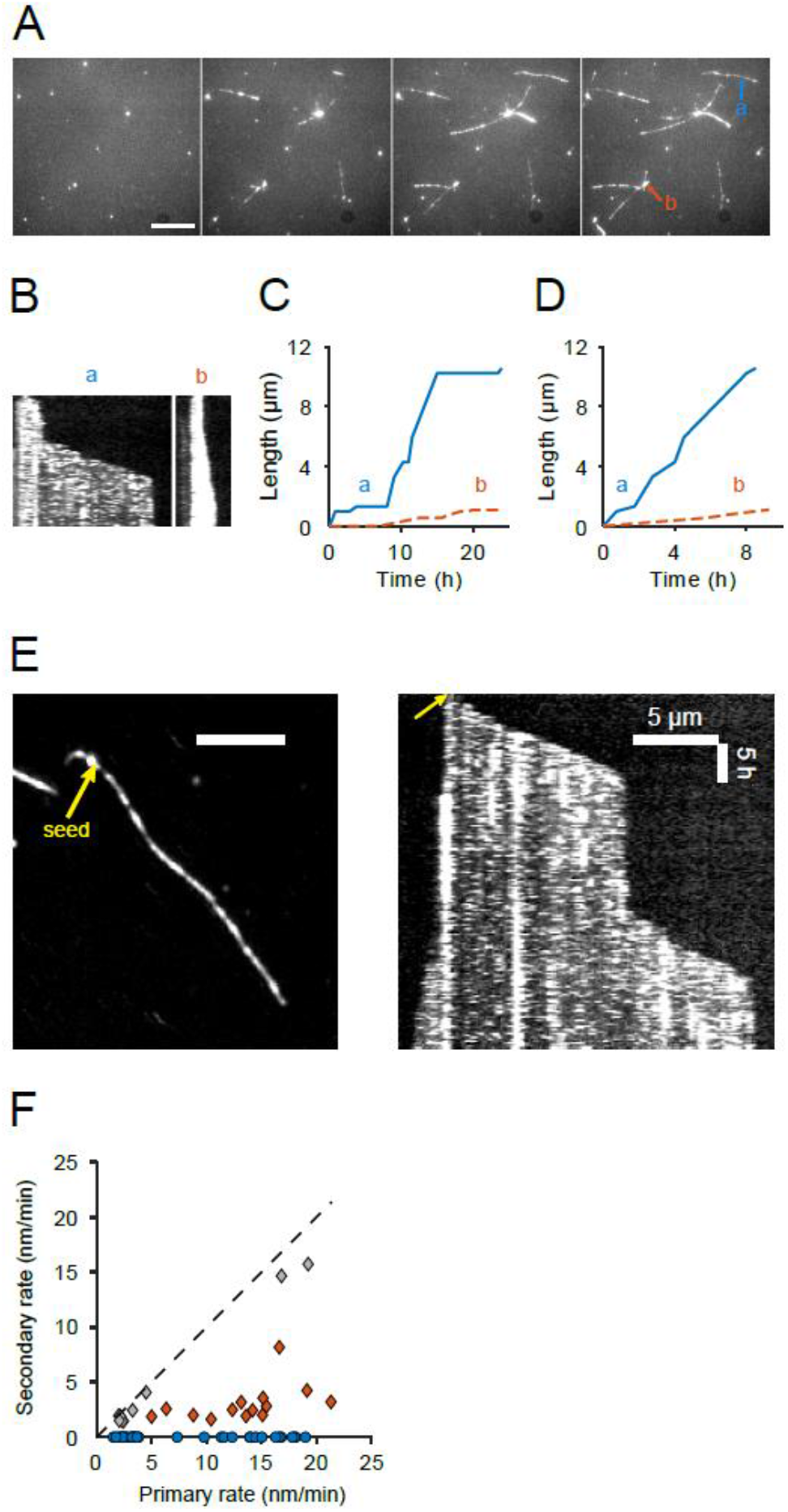
Real-time MoPrP 91 fibril elongation analysis observed by TIRFM. (A) a representative region imaged at multiple time points (0 h, 8 h, 16 h and 24 h, respectively) showing the elongation from dot-like seeds into long, isolated fibrils or fibril clusters (2 M GdnHCl, 1 μM MoPrP 91, 37 °C). The scale bar represents 10 μm. (B) kymographs of two fibrils (a) and (b) indicated by blue or orange arrows in the last panel of (A), respectively. The x-axis in each kymograph represents length along a line passing through the fibril axis, and y-axis corresponds to time. (C) length versus time traces of fibril a (blue solid line) and b (orange dashed line) converted from fibril edges in kymographs. (D) length versus time traces of fibrils a and b after removing stalls. (E) left panel: the image of a fibril elongated for 47 h, generated by running z-projection (projection type: standard deviation) command in ImageJ for a stack of 190 slices recording the growth of this fibril with 15 min interval. Arrow points to the original seed at t0. Right panel: kymograph of the fibril. (F) scatter plot of fibrils’ pause-free rates showing growth directionality. Blue, circular dots on x-axis represent pause-free rates of unidirectional fibrils; each diamond-shaped dot (orange or grey) represents a bidirectionally growing fibril with the pause-free rate of its faster end on x-axis and slower end on y-axis.

Fibril growth was interrupted by stall phases in a ‘stop-and-go’ growth. This growth pattern had previously been observed in other amyloid fibrils (Wordehoff et al., 2015; Young et al., 2017) and attributed to structurally incorrectly bound monomers/oligomers to the fibril ends, which have to be detached or covert to the correct structure before other molecules can add (Buell, 2019). We calculated the pause-free growth rates (Figure 1D) and relative stall time percentages for further analysis. Strikingly, growth rates were mostly constant for individual fibrils, however, they could vary dramatically between different fibrils, as shown for fibrils ‘a’ and ‘b’ in Figure 1A-D.

Fibril growth was highly directional (Figures 1A, E, F). Figure 1F plots pause-free rates of fast (forward) versus reverse growth in a scatter plot with each dot representing the growth pattern of a single fibril. Most fibrils grew in one direction only (blue dots) or showed reverse growth being much slower than forward elongation (red diamonds). These data suggest that fibrils are structurally distinct at opposing ends, leading to easier and faster incorporation of PrP at one end over the other. Indeed, recent atomic models of PrP amyloid fibrils (Wang et al., 2020) and prion rods (Kraus et al., 2021; Manka et al., 2022b) feature a step pattern at the fibril end, resulting in two asymmetric growth surfaces. A small portion of fibrils, marked by grey diamonds, elongated at similar rates in both directions. Since light microscopy cannot resolve single seeds and seed clusters, these fibrils were likely seeded by clusters that happened to be aligned in their fibril growth axes.

For a quantitative analysis of elongation kinetics, the growth of 114 fibrils was analyzed as described above (Figures 2A and 2B). Strikingly, despite all fibrils being seeded from the same seed population, growth rates did not follow a single distribution as had previously been observed for other amyloid fibrils (Young et al., 2017). Rather, growth rates could be grouped into two clusters featuring slow or fast growth and distinct fluorescence intensities.

**Figure 2.**
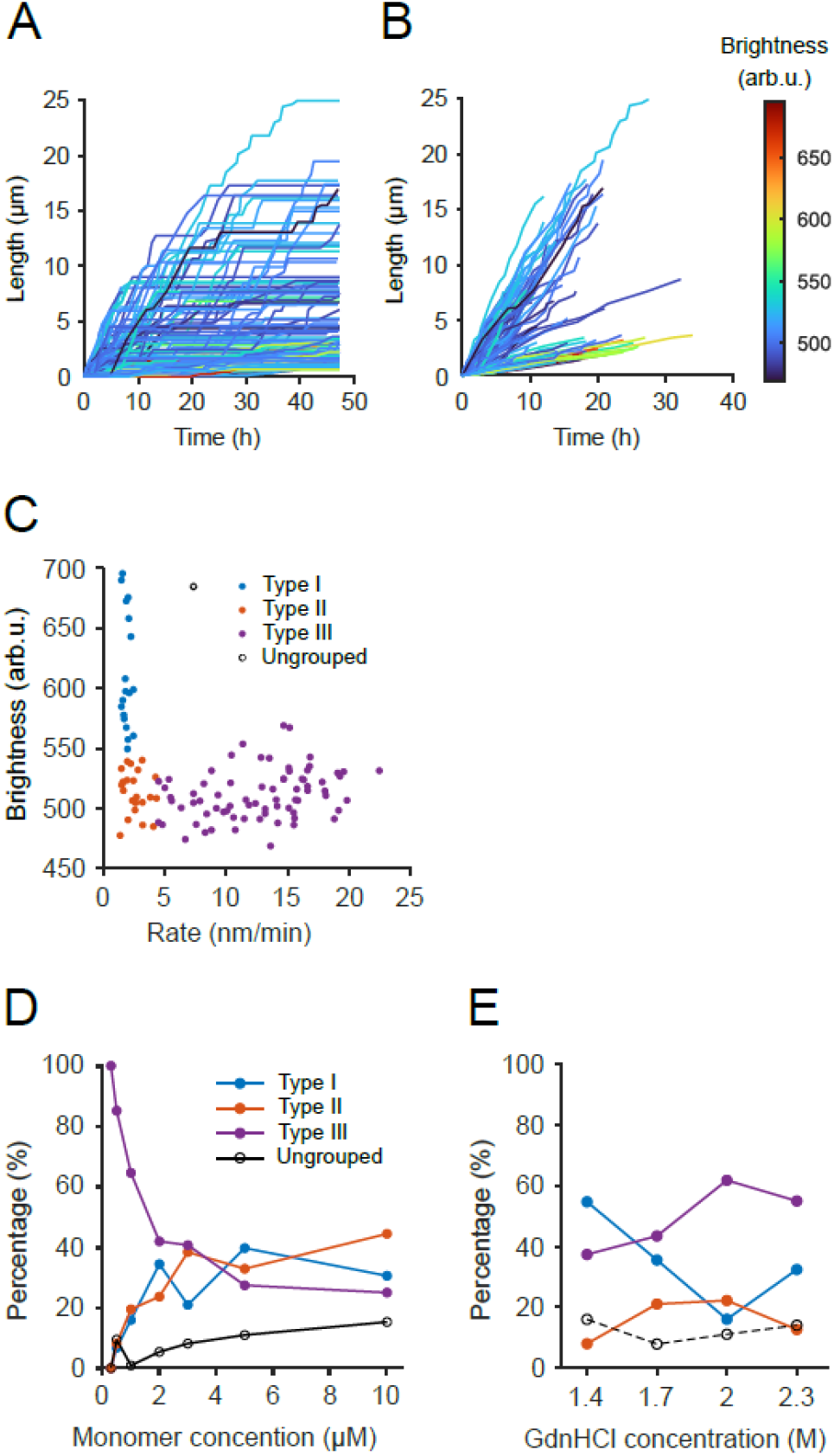
Kinetic analysis revealed multiple fibril types. (A) length versus time traces for fibrils elongated in 2 M GdnHCl with 1 μM PrP^C^ at 37°C. Traces are coloured coded by fibril brightness. (B) length versus time traces with stall phase removed. (C) scatter plot of fibril’s brightness versus pause-free rate. Fibrils were grouped based on the clustering within the scatter plot: slow-growing and bright fibrils as type I (blue), slow, dim fibrils as type II (orange), and fast, dim fibrils as type III (purple). (D) dependence of three fibril types’ fractions on PrP^C^ concentration. The total fibril number under each condition is the sum of type I, II and III fibrils, not including ungrouped fibrils. At 1 μM PrP^C^, fibrils in five FOVs are analysed, and standard deviations of fractions among those five FOVs are 11.5%, 12.7% and 13.7% for type I, II and III fibrils, respectively. (E) dependence of fibril fractions on GdnHCl concentration.

Growth traces in Figures 2A and 2B are colour-coded by fibril brightness, showing that most slow-growing fibrils were brighter than fast fibrils. A scatter plot of fibril brightness versus pause-free rate (Figure 2C) revealed the presence of three distinct types of fibrils: slow-growing bright fibrils (denoted as type I fibrils, *blue*), slow-growing dim fibrils (type II, *red*), and fast-growing dim fibrils (type III, *purple*), while a small portion did not fall into any group or displayed inconsistent growth rates (open circles). Thus, bright fibrils were exclusively slow-growing and all fast fibrils were relatively dim. Those differences likely represented different fibril structures as will be discussed below.

This methodology then allowed us to analyze how monomer concentration, degree of folding and temperature determined stall percentage, growth rate and partitioning between different types of fibrils in a large data set.

### Elongation of competing PrP fibril populations

Monomer concentration determined relative populations of type I, II and III fibrils (Figure 2D). Type III fibrils were favoured at lower monomer concentrations, accounting for over 80% below 1 μM, while high PrP concentrations shifted the equilibrium towards slower-growing type I and II fibrils.

To test how the unfolding of substrate PrP affected the fibril populations, elongation experiments were performed at GdnHCl concentrations ranging from 1.4 M to 2.3 M, corresponding to a fraction of 0.8% – 52% unfolded protein under assay conditions (Figure S1C, D). Figure 2E shows that low denaturant concentrations favoured the formation of bright type I fibrils. Their population decreased with GdnHCl concentration, while the population of fast-growing type III fibrils increased. It should be noted that fast-growing fibrils occasionally detached from the surface at 2.3 M GdnHCl. Those fibrils were eliminated from rate analysis, which artificially depressed the fraction of type III fibrils. To summarize, an increasing fraction of unfolded PrP monomer shifted the equilibrium between competing fibril seeds towards fast-growing, type III fibrils.

### PrP fibrils with different kinetic profiles correspond to distinct structures

Next, we tested whether fibril types corresponded to distinct fibril structures by choosing reaction conditions, which favoured the growth of type I or type III fibrils, respectively: (a) 1 M GdnHCl with 1 μM monomer (~ 70% of type I fibrils) and (b) 2 M GdnHCl with 1 μM monomer (~ 60% of type III).

To visualize fibril structures, we performed elongation experiments *in situ* on electron microscopy grids, imaged the elongated fibrils by EM and analyzed fibril diameters under both conditions (Figures 3A and 3B). Since it is difficult to identify newly grown fibrils with certainty in this type of experiment, we only analyzed fibril segments that were longer than 500 nm, which was much longer than the typical length of the initial seed fibrils (Figure S1B). PrP fibrils grown under the two conditions had distinct distributions of fibril widths: 16 - 18 nm wide fibrils were dominant in condition (a) favouring type I fibrils, while condition (b) favouring type III fibrils generated mostly 6 - 8 nm wide fibrils. As can be seen in Figures 3A and 3B, the two different fibril widths likely represent double-strand and single-strand fibrils, respectively. Single-strand fibrils observed under condition (b) tended to be much longer than fibrils under condition (a) reflecting faster fibril growth.

**Figure 3.**
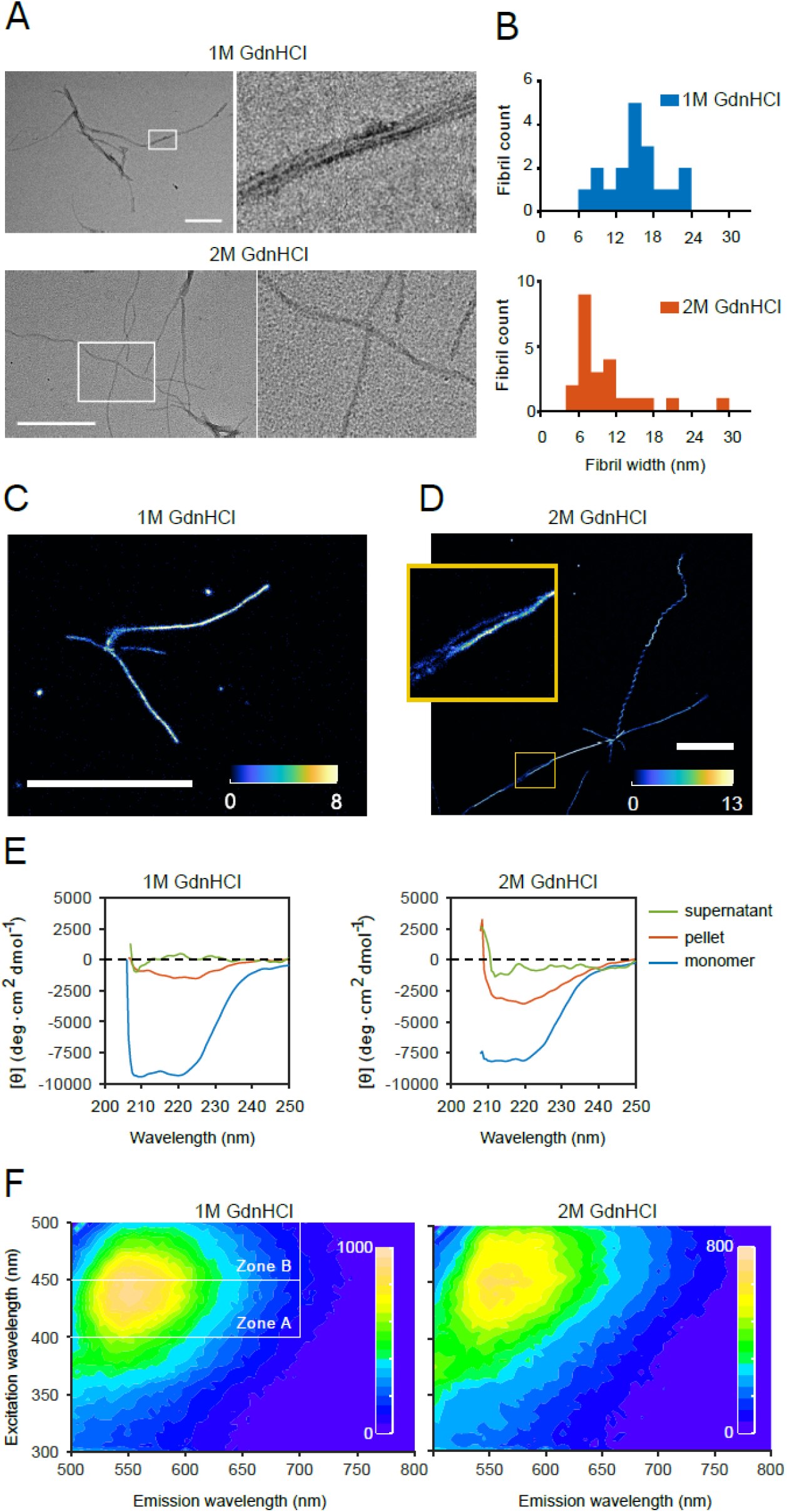
PrP fibrils with different kinetic profiles correspond to distinct structures. (A) typical TEM images of fibrils elongated in 1 M GdnHCl (top) or 2 M GdnHCl (bottom). Scale bar 500 nm. (B) width distribution of fibrils grown in 1 M GdnHCl (top) or 2 M GdnHCl (bottom) (C) typical TAB image of fibrils elongated in 1 M GdnHCl. Images were acquired in the presence of 50 nM Nile Red. Brightness represents the number of localizations identified in each pixel. Scale bar 5 μm. (D) TAB image of fibrils elongated in 2 M GdnHCl and a zoomed-in view showing the branching of a fibril. (E) CD spectra of PrP^C^ or PrP aggregated in buffer with 1 M (left) or 2 M (right) GdnHCl. For aggregated PrP, ultracentrifugation was performed; pellet resuspension and supernatant were measured by CD separately. (E) exemplary contour maps for samples generated by sequential seeding in buffer containing 1 M GdnHCl (left) or 2 M GdnHCl (right). The partition of zones A and B is shown in the left panel.

To confirm this conclusion, fibrils grown under conditions (a) and (b) were analyzed by transient amyloid binding (TAB) microscopy (Figures 3C and D). The colours represent the number of individual dye molecule localizations, i.e. ‘brightness’. While amyloid fibrils are too narrow to directly resolve single and double-strand fibrils, two distinct populations of fibrils are visible: short, straight bright (type I) fibrils, which occasionally unspliced (Figure 3D, inset) revealing their multi-strand structure. In contrast, TAB and EM both resolved single strands with mostly twisted morphologies for type III fibrils. Interestingly, both EM and TAB imaging identified additional fibril morphologies, albeit at low abundance. We observed single-strand straight fibrils, single-strand curved fibrils, multiple-strand fibrils with regular twisting (Figure S2B), multiple-strand fibrils without twisting and helically coiled fibrils (Figure S2A-C). This variety of fibril morphologies suggests that PrP fibrils, rather than being a single, homogenous species, can exist in multiple amyloid structures, which likely form from subtly different underlying folds of the peptide chain. These fibril polymorphs were missed by the lack of spatial resolution in TIRF microscopy.

EM and TAB images suggest that fibril types may not only differ in their number of strands but also in their internal structures. To substantiate this hypothesis, the secondary structures of fibrils formed under both conditions were studied by CD spectroscopy (Fig 3E). Two rounds of seeding in solution were performed for conditions (a) and (b) condition to enrich type I and type III fibrils, respectively. Under both conditions, the spectra of the pelleted and resuspended fibrils had minima between 218 – 225 nm indicating the presence of β-sheets. However, spectra were distinct with minima at ~220 for type III, and at ~225 nm for type I enriched fibrils respectively, confirming that fibrils generated under the two conditions had distinct secondary structures.

This interpretation was confirmed by spectral fingerprinting of the luminescent conjugated oligothiophene dye (LCO) dye heptamer formyl thiophene acetic acid (h-FTAA). LCO dyes display spectral shifts, which can differentiate structural subtypes of amyloid (Aslund et al., 2009). Fig. 3F shows 3D excitation-emission contour plots of h-FTAA bound to PrP fibrils grown under conditions (a) and (b) from seed also made under the same condition. These fibrils are denoted ‘aa’ and ‘bb’, respectively, in Figure 4A. Their spectra differ both in peak shape as defined by the ratio of fluorescence emission integrated over zones A and B and in their maximal fluorescence intensities (Figures 3E and 4A). Both, CD spectroscopy and spectral fingerprinting, therefore, support that fibril types I and III not only correspond to double- and single-strand fibrils, respectively, but that both types of fibril feature different internal structures. This result suggests distinct assembly mechanisms that propagate their structural characteristics.

**Figure 4.**
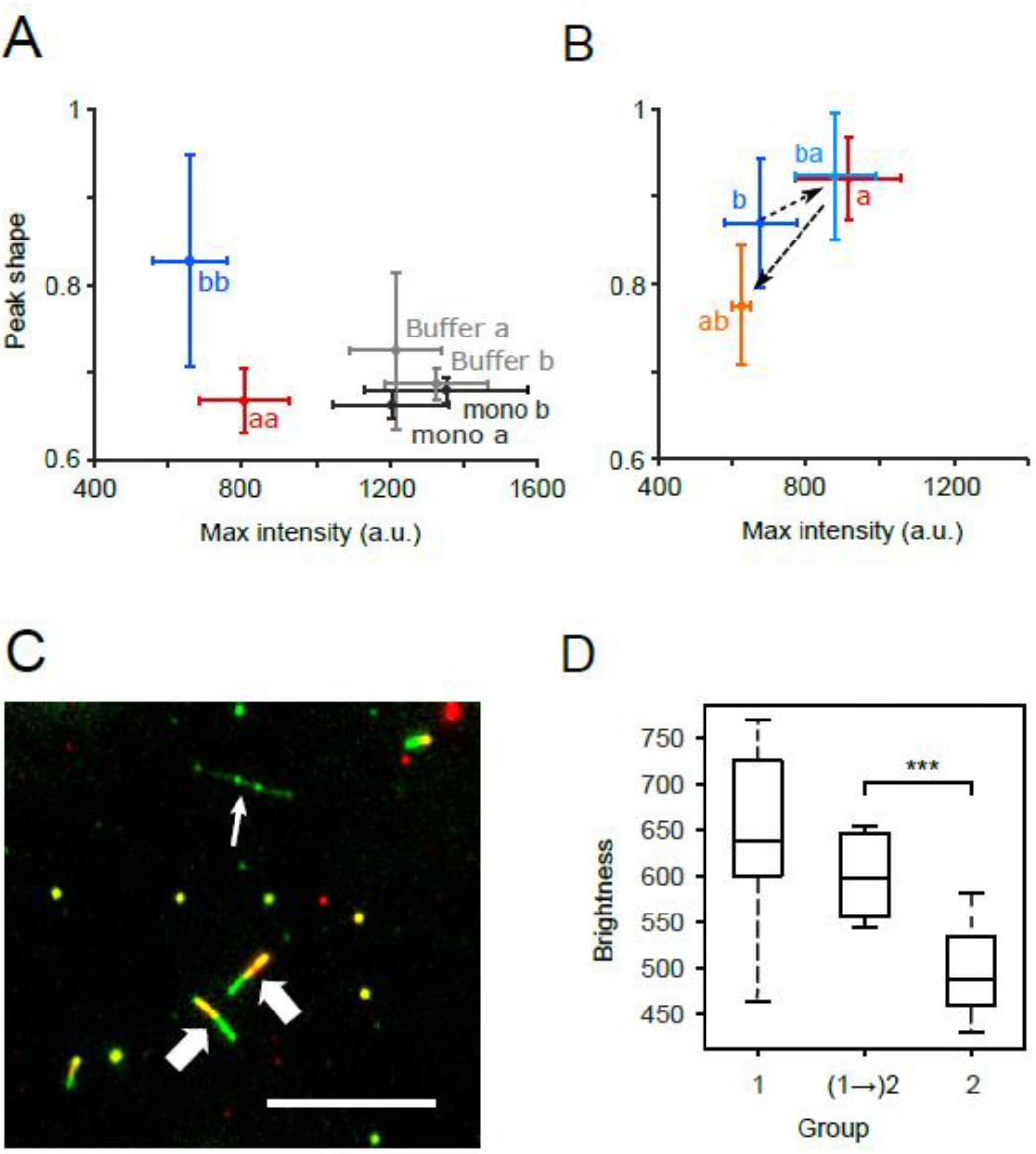
Fibril types propagate faithfully under competing conditions. (A) the plot of peak shape (integral of zone B/integral of zone A) against maximum intensity for samples aa and bb which were seeded twice in the corresponding buffer, and for buffer and monomer samples in the corresponding buffer a and b. (B) comparison of peak shape versus maximum intensity for samples a, ab, b and ba. (C) composite image of an ROI showing fibrils which have elongated in buffer with 1 M GdnHCl for 71 h (green channel), and fibrils after a sequential seeding in 2 M GdnHCl (red channel) for 41.5 h. PrPC concentration was 1 μM in both seeding phases. The thin arrow refers to a fibril that only grew in the second assay; fat arrows point to fibrils which elongated in both seeding phases. The scale bar represents 10 μm. (D) brightness analysis for fibril segments that grew in the first seeding phase (group 1), fibril segments that grew in the second seeding phase from an elongated fibril in the first phase (group (1 →)2), and fibrils that exclusively grew in the second phase (group 2). 19 fibrils in total were analysed.

### Fibril types retain structural characteristics under adverse growth conditions

We next determined whether fibril types could faithfully propagate their respective structures under adverse conditions favouring a different fibril type or whether fibril types were determined by solution conditions. Seeds grown in condition (a) favouring type I fibrils were elongated in condition (b) favouring type III fibrils and vice versa. When fibrils were grown in solution and analysed in bulk, second-generation fibrils grown in condition (a) (Figure 4B; ‘ba’) had similar spectral fingerprints to first-generation seeds ‘a’. Likewise, the footprint of first-generation ‘a’ seeds shifted when elongated under (b) conditions (Figure 4B, ‘ab’). This result seems to suggest that solution conditions rather than templating determined fibril structure. However, a closer analysis of single fibril growth revealed this not to be the case.

To probe templating fidelity in fibril growth, we designed a two-phase experiment. In the first phase fibrils grew from seeds under buffer conditions (a) favouring the formation of type I fibrils. Then, the buffer was switched to (b) and fibril growth was monitored for a further 88 h by TIRF microscopy. In Figure 4C, fibrils present at the beginning of the second phase are shown in red, and fibrils grown or elongated during the second phase are shown in green. Correspondingly, fibrils from the first phase appear yellow, if still present in the second phase of the experiment, while red-stained dots correspond to initial seeds that had detached during the second phase.

In the second elongation phase, type I fibrils continued to grow (Fig. 4C bold arrows); type III fibrils started to grow from dot-like seeds that hadn’t extended in phase 1 or at a position where no obvious seed was observed (Fig 4B, thin arrow). Figure 4D plots the brightness profiles of fibril segments grown in the first and second phases. Fibril brightness did not significantly change on elongation after the buffer exchange (Figure 4D, 1→2). These data indicate that elongating fibrils retained their fibril structures after the switch of buffer conditions and that type III fibrils were not cross-seeded by type I and vice versa. Our data, therefore, suggest that buffer conditions shift the equilibrium between fibril types, so that type I fibrils could outcompete type III in condition (a) and vice versa, as shown in Figure 4B, but that type and structure of every single fibril was specifically templated by the seed type rather than determined by buffer conditions.

### Competing PrP fibril populations elongate with distinct mechanisms

Specific templating of fibril types suggests the presence of distinct elongation mechanisms. To test this hypothesis, we analyzed the dependence of fibril elongation on PrP monomer concentration, denaturant concentration and temperature. Figure 5A and Figure S3B, C show the distributions of stall-free growth rates and stall percentages for fibrils grown at 0.3, 0.5, 1, 2, 3, 5 and 10 μM monomer concentration, respectively, separated by fibril types, while Figure S3A represents the data for the entire fibril population. The three fibril types exhibited starkly different concentration dependencies of their pause-free elongation rates (Figure 5B) and stall percentages (Figure 5C). These data suggest that different steps of the assembly mechanism were rate-limiting for different fibril types.

**Figure 5.**
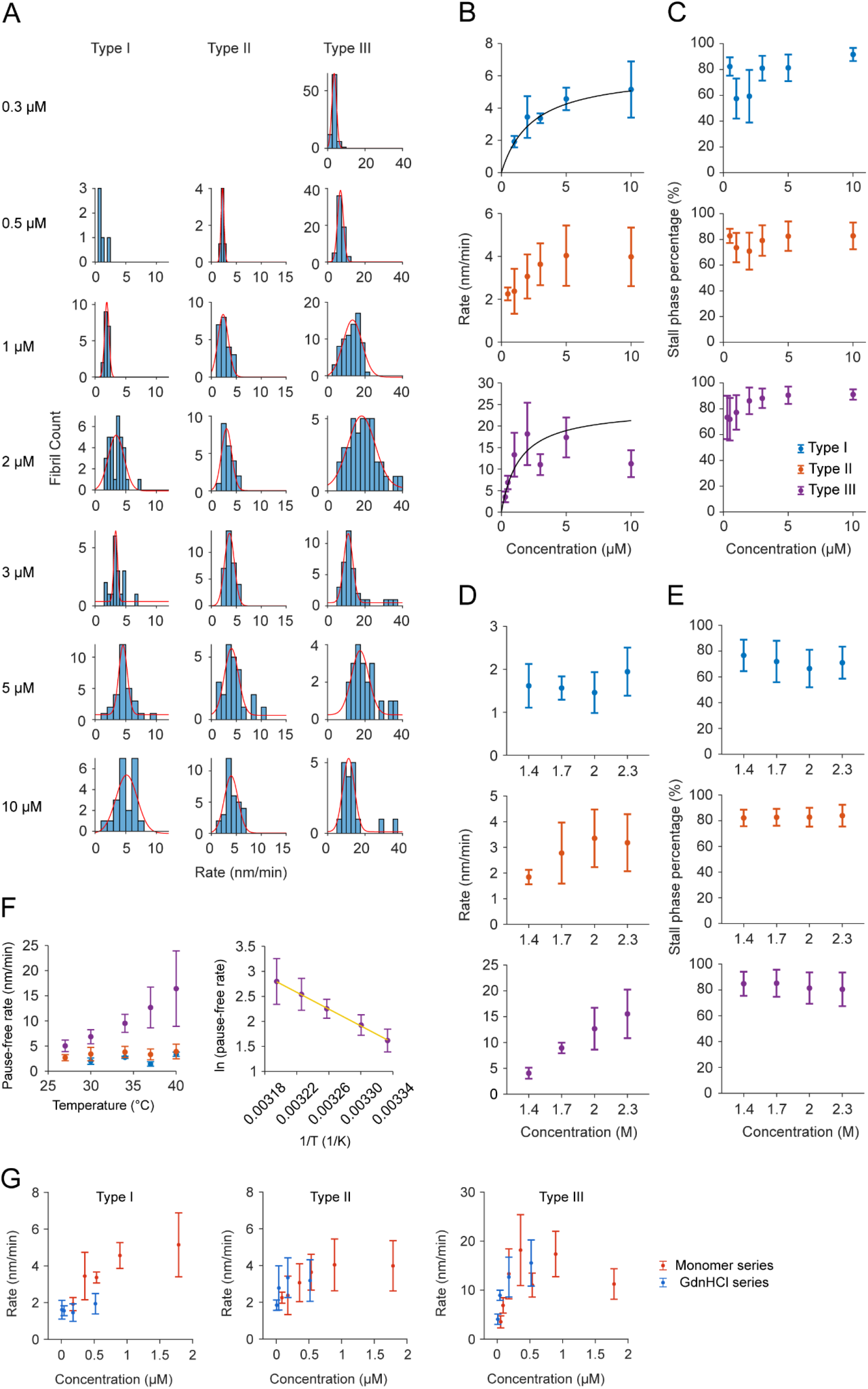
Kinetic analysis of individual fibril types under different conditions. (A) pause-free rate distribution of 3 types of fibrils in monomer concentration series. Red curves represent Gaussian fitting to the distribution. (B) dependence of pause-free rates of type I (top), type II (middle) or type III (bottom) fibrils on PrP^C^ concentration. Pause-free rates are the peak positions taken from Gaussian fittings in (A), and error bars represent σ. The dashed curves in the top and bottom panels represent a Michaelis-Menten fitting to data. In the bottom panel data points at 3 μM and 10 μM were removed from the fitting. (C) dependence of stall phase percentages of type I, II or III fibrils on PrPC concentration. Stall percentages are the average value of fibrils of the specific type. Errors are standard deviations. (D) dependence of pause-free rates of type I (top), II (middle) or type III fibrils (bottom) on GdnHCl concentration. (E) dependence of stall phase percentages of type I, II or III fibrils on GdnHCl concentration. (F) temperature dependence of fibril pause-free rate. Left: pause-free rates of type I (blue), II (orange), or III (purple) fibrils under different temperatures. Right: Arrhenius plot for type III fibrils. The straight line is a linear fit of the five data points. (G) Dependence of pause-free rate of three fibril types on unfolded monomer concentration in PrP^C^ concentration series (red) and GdnHCl concentration series (blue). See also Figures S3-S5 for fibril distribution histograms.

Both type I and III fibril growth accelerated with increasing monomer concentration. Fast-growing type III fibrils saw a strong linear concentration dependence between 0.3 – 2 μM, with a maximal rate of 20 ± 6 nm/min, while higher PrP concentrations inhibited fibril growth. In contrast, bright, slow type I fibrils exhibited a weaker concentration dependence and did not show inhibition at high PrP concentrations.

In the simplest case, fibril elongation could be described as catalytic conversion of monomer (M) to fibril units (F) at the end of a growing fibril (E) in which the successful conversion of the monomer yields a fresh catalytic interface for further fibril growth (Eq 1). This situation is analogous to a Michaelis-Menten type reaction scheme (Eq 2).

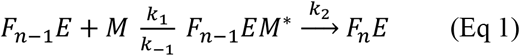

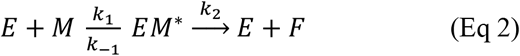

We have analyzed elongation rates under the assumption of excess monomer (steady-state) to calculate K_m_ = (k_-1_ + k_2_) /k_1_ and k_2_, respectively (line graphs in Fig 4B). This simple model fits the concentration dependence of type I fibrils with K_m_ = 2.4 ± 0.3 μM, and k_2_ = 0.44 ± 0.05 s^-1^, assuming type I fibrils were made of double strands. It could adequately describe type III fibril growth at low monomer concentrations (K_m_ = 1.4 ± 0.5 μM, k_2_ = 0.8 ± 0.2 s^-1^), but failed to capture the inhibition of type III fibril growth at higher monomer concentrations. Non-competitive inhibition by unproductive binding of PrP monomers could account for this inhibition (Honda and Kuwata, 2017). Alternatively, PrP could be forming off-pathway assemblies at high protein concentrations, which bind at the fibril end and prevent fibril growth.

In contrast to types I and III, type II fibril growth showed a negligible concentration dependence overall, suggesting that the rate-limiting step of elongation was 0th order with respect to PrP concentration. This behaviour is compatible with the structural conversion of PrP being rate-limiting.

To further analyze the assembly mechanism, we determined the dependence of fibril growth rates on denaturant concentration from 1.4 – 2.3 M GdnHCl at a fixed PrP concentration (Figures 5D, E and S4). Under our reaction conditions, elongation rates of type III fibrils increased linearly with unfolded PrP concentration, suggesting that this fibril type grows by incorporating unfolded PrP molecules. Indeed, data from GdnHCl and monomer concentration series fell into one graph when plotting elongation rate against unfolded monomer concentration (Figure 5G). In contrast, type I and II fibril elongation showed no significant dependence on GdnHCl concentration. This behaviour is compatible with the incorporation of a partially folded intermediate, whose concentration changes weakly in the GdnHCl range probed in the experiment.

Stall percentages increased with PrP monomer concentration for all three fibril types (Figures 5C and S3B). While stall percentage was low (57 ± 16%) for type I fibrils at 1 μM PrP, it increased markedly to 91 ± 5% at 10 μM monomer concentration. Type II and III fibril stalling increased from 73 ± 11% to 86 ± 7% and 76 ± 12% to 85 ± 10%, respectively, indicating that stall events dominated at high PrP concentrations. Both, fibril stalling and the drop in elongation rate at high monomer concentrations observed for type III fibrils, indicate a concentration-dependent negative feedback loop, which inhibits fibril growth. While both processes appear to operate in different time scales, the distinction is somewhat arbitrary, because it depends on the sampling time of our kinetic assay with stall events < 15 min being perceived as a reduction in elongation rate. Both stalling and slowed elongation are most likely due to PrP bound to the fibril ends, either in the form of monomers or non-fibrillar assemblies, which could not convert into the amyloid fold. It is possible that, despite the presence of GdnHCl, high PrP concentrations facilitated the formation of non-fibrillar PrP assemblies that could contribute to the stalling of fibril growth. However, GdnHCl concentration did not change the stall percentages for any of the three fibril types, arguing against this interpretation.

Elongation rates were measured at five temperatures (27, 30, 34, 37, and 40°C) at fixed solution conditions (2 M GdnHCl, 1 μM PrP^C^) to determine the activation energies for elongation of the three fibril types (Figure 5F and S5). The fraction of unfolded protein varied from 5% at 27°C to 40% at 40°C under these conditions (Figure S1E, F). Pause-free elongation rates for type III fibrils had a strong temperature dependence, which followed the Arrhenius equation with activation energy (E_a_) of 70 ± 2 kJ/mol. This is comparable to the literature value of ~ 50 kJ/mol under conditions when unfolded monomer is abundant (Milto et al., 2014). In contrast, pause-free rates for type I and II fibrils did not change with temperature, suggesting the rate-limiting step was not temperature-dependent. This may be due to entropic and enthalpic contributions to the activation energy largely cancelling each other out (Buell et al., 2012).

In summary, our single-fibril kinetic analysis determined that at least three different PrP fibril types were growing in competition from a seemingly homogenous pool of seeds. While growth conditions could shift the equilibrium between fibril types, fibrils elongated faithfully without apparent cross-seeding by distinct mechanisms, similar to what has been posited for prion strains. Our data suggest the presence of a polymorphic ‘cloud’ of fibril conformations, which compete for the same substrate pool, raising the question of whether authentic prion strain elongation shares the same structural and mechanistic diversity.

### Elongation of authentic prion rods

We analyzed the elongation of two mouse prion strains, RML and ME7, in the presence of recombinant monomer to probe how their elongation kinetics and fibril structures differ from synthetic fibril seeds and between different prion strains.

RML and ME7 prion rods were purified from infected mouse brain as previously described (Terry et al., 2019; Wenborn et al., 2015), adsorbed onto coverslips and incubated with MoPrP 91 monomer (10 μM) in 2M GdnHCl. RML seeds readily templated fibril growth under these conditions (Figure 6A). In contrast, only few short fibrils grew from ME7 seeds. A quantitative analysis was performed for N = 12 RML-seeded fibrils, and the growth traces and stall percentages are shown in Figure 6B and C, respectively. Similar to the fibrils grown from synthetic seeds, pause-free rates within a fibril were nearly constant. However, fibrils displayed diversity in growth rates, stall percentages and fibril brightness (Fig 6B, C). Elongation rates of RML seed (0.4 - 1.7 nm/min) were generally slower than all three types of synthetic seeds grown under the same condition (Fig 6D). However, unlike recombinant seed elongation, rates and brightness of RML seeded fibrils did not correlate, so no distinct fibril types could be identified.

**Figure 6.**
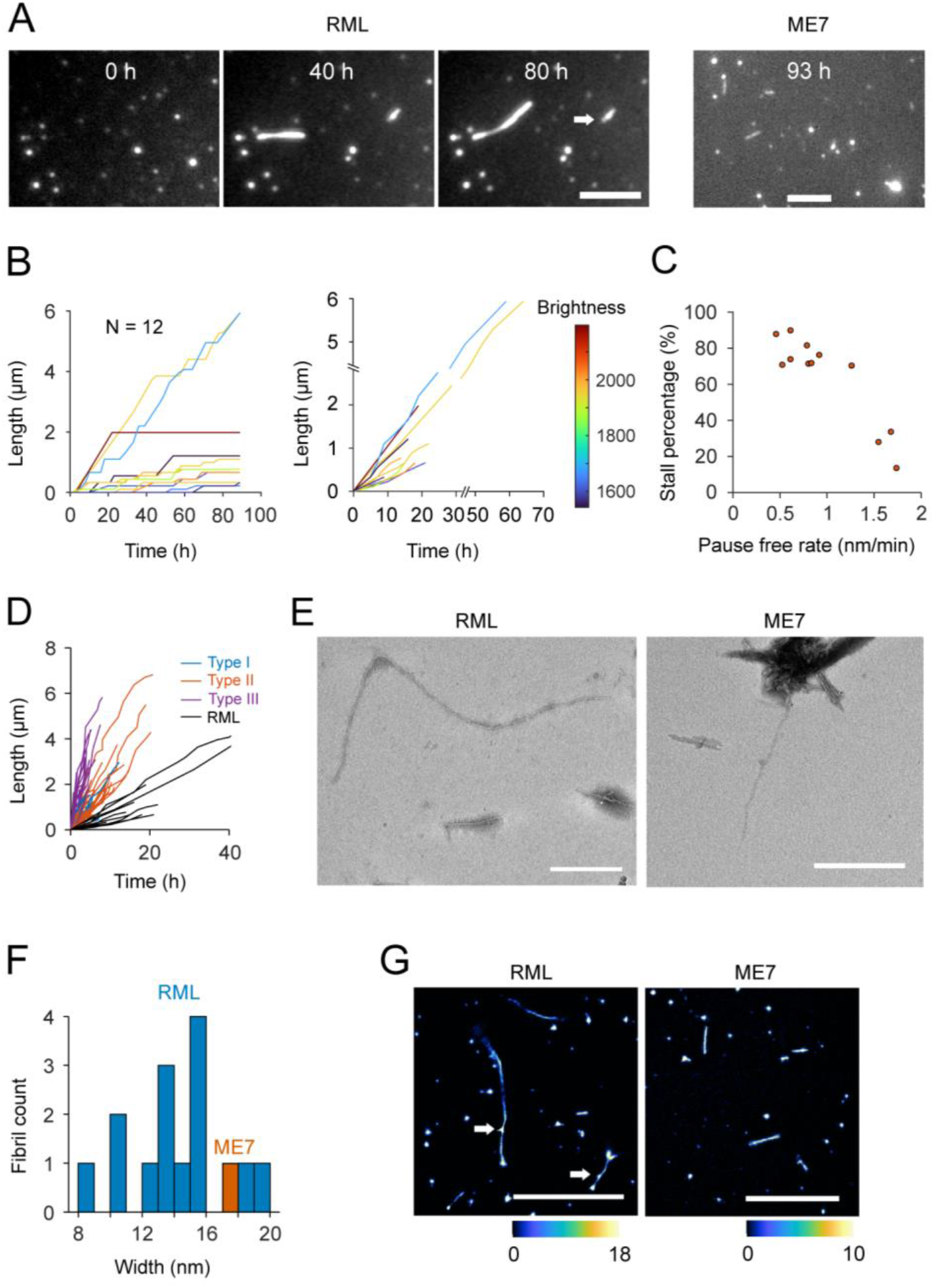
Elongation of authentic rods. (A) TIRFM images showing the elongation of RML and end point of ME7 fibrils. The scale bar represents 5 μm. (B) length versus time traces for 12 RML fibrils with stall phases included (left) or removed (right). Traces were colour coded by fibril brightness. (C) stall percentages of each fibril plotted against the brightness. Stall percentage was calculated using the time before the last stall phase as total time. (D) length versus time traces with stall removed of RML overlaid with grouped elongation traces of recombinant seeds. (E) TEM images of RML (left) or ME7 (right) fibrils elongated on EM grids. The scale bar represents 500 nm. (F) width distribution of RML fibrils and an ME7 fibril elongated on EM grids. (G) TAB images of elongated RML fibrils. Scale bar 5 μm. Arrows in panels A and G indicate bidirectional growth.

*In situ* elongation experiments with RML and ME7 prion rods on EM grids confirmed that RML prion rods readily elongated into fibrils with a mostly twisted architecture (Figure 6E) and an average width of 12-16 nm (Figure 6F). In contrast, the few fibrils that could clearly be identified as having been newly seeded from ME7 prion rods were short, single-strand fibrils with a diameter of ~18 nm. Similar to EM, TAB microscopy showed long, often bent fibrils growing from RML seeds, while ME7 yielded short, straight fibrils (Figure 6G). Intriguingly, fibrils grown from RML and ME7 prion rods displayed repeating intensity patterns with a period of 134 ± 4 nm and 153 ± 8 nm, respectively (Figure S6). These periods coincide with the crossover distances of RML and ME7 prion fibrils recently determined by cryo-EM (Kraus et al., 2021; Manka et al., 2022a; Manka et al., 2022b), suggesting that elongation of prion rods preserves elements of prion fibril architecture.

Overall, our data suggest that both RML and ME7 rods are capable to elongate under the conditions examined here, but that growth rates are starkly different between the two strains. The elongation rates of RML-seeded fibrils are slow compared to authentic fibrils. A diversity of fibril brightness and elongation rates suggest that the purified RML prion samples can template more than one fibril structure. Whether this points to a lack of fidelity in templating their structure under denaturing conditions or whether it indicates the presence of structural heterogeneity of a prion quasispecies will have to be resolved by future kinetic studies under physiological replication conditions.

## Discussion

Unique fibril structures, which can be replicated by templated monomer addition are thought to be the structural basis for prion strains. Cryo-EM of three prion strains, RML (Manka et al., 2022b), ME7 (Manka et al., 2022a) and 263K (Kraus et al., 2021), albeit propagated in two different species, displayed homogenous structures with similarities in their overall arrangement of the polypeptide chain. However, their subtle differences in local fold yielded massive changes in overall fibril morphology. Distinct structures of patient-derived amyloid corresponded to specific tauopathy phenotypes (Falcon et al., 2018a; Falcon et al., 2018b; Fitzpatrick et al., 2017; Scheres et al., 2020). Rapid progressing AD seeded distinct fibril structures from normal sporadic AD (Qiang et al., 2017) and Aβ42 fibrils that were isolated from patients suffering from sporadic and familial AD showed two distinct morphologies (Yang et al., 2022). Notably, fibril structures in each patient, while not entirely homogenous, were dominated by one isomorph, suggesting competing templating processes within the affected patient brain.

The environment can steer fibril formation towards amyloid with different structures and stabilities. Aβ 40 can form multiple stable amyloid structures *in vitro*, whose populations depend on metal ions, pH, temperature and the presence of cofactors (Kodali et al., 2010; Niu et al., 2014; Petkova et al., 2005) and salt concentration directs α-synuclein into two distinct fibril morphologies, which can propagate faithfully (Bousset et al., 2013). Similarly, polymorphs of SAA can template distinct fibril structures (Bansal et al., 2021; Heerde et al., 2022). Full-length hamster PrP could form two distinct fibril types under different shaking modes (Makarava and Baskakov, 2008), which differed in their morphologies, internal structures and ThT fluorescence. The unique fibril type was maintained after seeding even under unfavourable shaking modes. Our single-particle kinetic analysis revealed the presence of structurally distinct PrP fibrils that coexisted *in vitro* and faithfully templated their structures during elongation, while at the same time competing for monomer addition (Figure 4). The energy barrier for monomers adopting an existing seed structure has a low free energy barrier, compared to a change in structure, leading to faithful elongation, even though the seed structure might not be the most stable one under the specific condition (Hadi Alijanvand et al., 2021).

Structural polymorphs were also observed within homogenous PrP amyloid preparations (Aubrey et al., 2020; Goldsbury et al., 2005). *De novo* aggregation of hamster PrP 90-231 (Alvarez-Martinez et al., 2011) yielded distinct aggregation kinetics and morphologically different fibrils under the same conditions, which suggested the formation of structurally different nuclei. However, studies of fibril formation in bulk are limited in their resolution of competing species within a homogenous assay and fail to resolve how the structures of individual fibrils correlate with their elongation dynamics and mechanisms.

Single-molecule measurements on PrP seeded elongation using TIRFM revealed three main groups of fibrils based on their brightness and kinetic profile (Figure 2C). In comparison to high-resolution TEM images of fibrils grown on EM grids (Figures 3A and 3B), the fast-growing, dim fibrils in TIRF measurements were likely single-strand fibrils, while the slow-growing and bright fibrils likely corresponded to double-strand fibrils. Thus, fibrils displayed distinct dynamic properties, which were linked to their specific structures. Interestingly, fibrils seeded by RML and ME7 prion rods all had diameters similar to the double-stranded type I fibrils, elongated more slowly and lacked the dynamic heterogeneity of fibrils seeded from synthetic seeds (Figure 6). This observation may reflect a greater degree of conformational restraint imposed by the prion seed corresponding to a lack of structural diversity in prions when compared to PrP amyloid fibrils (Kraus et al., 2021; Manka et al., 2022b; Wang et al., 2020). It is tempting to speculate that PrP amyloid growth may compete with prion replication in vivo and that the kinetics favour amyloid formation, while templating of prions produces more stable but slower growing fibrils.

In our assay, elongation was the only process observed for pre-existing seeds in our surface-based elongation experiment. Fibril elongation was also dominant in previous studies of multiple amyloidogenic proteins, such as Aβ40 (Ban and Goto, 2006), Aβ42 (Young et al., 2017), α-Syn (Wordehoff et al., 2015) and Sup35 (Collins et al., 2004; Konno et al., 2020), but in these systems, no competing fibril species were observed. Under our conditions, we observed no fibril branching (Andersen et al., 2009) or fragmentation (Sang et al., 2018). This suggests that our reaction conditions did not favour secondary nucleation or fragmentation, simplifying the kinetic analysis.

Elongation of all three fibril types followed a ‘stop-and-go’ pattern over multiple time scales (Figure 2A), as observed for other protein amyloids (Wordehoff et al., 2015; Young et al., 2017). Intermittent growth suggested binding of PrP species that were not able to elongate, or trapping of the bound monomer in an incorrect conformation (Buell, 2019). While we cannot exclude that the glass surface contributed to fibril stalling, it is unlikely to be the dominant cause, because stall percentages depended on monomer concentration and solution conditions, whereas stalling due to surface interaction and steric hindrance would be expected to be independent of PrP concentration. Similarly, differences in fibril growth are unlikely to be caused by fibril orientation on the surface. If this were the case, we would expect ratios of type I, II, and III fibrils to be independent of monomer concentration and solution condition, which is contrary to our observations.

Similar to other amyloid fibrils (Young et al., 2017), elongation of synthetic PrP seeds was highly directional. The surfaces of the two fibril ends of a single are not symmetric (Kraus et al., 2021; Manka et al., 2022a; Manka et al., 2022b) which likely results in a slow and a fast interface for fibril growth. It is possible that in bidirectional fibrils type II represented the slow end and type III the fast-growing end of the same fibril. Our data suggest that RML seeds templated two different fibril types, representing fast and continuous growing fibrils and relatively slow fibrils which had longer stalls (Figure 6), which may correspond to single and double-strand fibril elongation, respectively. Cryo-EM revealed that 10% of RML prion fibrils were double-stranded (Manka et al., 2022b). Unlike most amyloid, these had mirror symmetry, indicating an anti-parallel orientation of strands and symmetric fibril ends [6]. Notably, bidirectional fibril growth was observed in some RML-seeded fibrils (Figure 6A and G, *arrows*), which could reflect this seed structure.

Different brightness and growth rates corresponded not only to single and double-strand fibrils but also to different assembly mechanisms. Low PrP concentration favoured type III fibrils; no type I fibril formation was observed below 0.5 μM, which could mean that type I fibrils could have higher critical concentration. Alternatively, monomer addition to double-stranded type I fibrils could follow higher reaction order kinetics than type III, whose growth was described by simple Michaelis–Menten -type kinetics. However, the low concentration dependence of type I growth argues against a higher-order mechanism.

Growth data are compatible with the recruitment of unfolded monomers into type III fibrils, while type I fibrils possibly recruited native-state monomers or an early unfolding intermediate. For type III fibrils, growth data of GdnHCl unfolding experiments could be mapped onto the concentration dependence data when correction for the degree of unfolding while type I fibril growth did not have a strong dependence on GdnHCl concentration (Figure 5G). Concentration dependence revealed a Michaelis-Menten type kinetics between 0.5 – 2 μM monomer concentration (Figure 5B) for type III fibrils, suggesting a first-order kinetics with respect to the precursor at low monomer concentration, while structural conversion became rate-limiting at higher PrP concentration. In addition, we observed a reduction in type III growth rate at PrP concentrations > 5 μM, which suggests an inhibition of fibril growth by natively folded monomer, according to the model formulated by Honda et al. (Honda and Kuwata, 2017). Temperature dependence of type III fibril growth followed Arrhenius’ law. The activation energy of 70 ± 2 kJ/mol is consistent with values calculated for the elongation of Mo PrP (89-230) fibrils in bulk seeding assays (Milto et al., 2014), with Ea ~170 kJ/mol under conditions where native-state PrP dominated, and Ea ~ 50 kJ/mol when PrP was predominantly unfolded.

## Conclusions

Our analysis revealed that, contrary to the long-time assumptions on amyloid growth, polymorphic populations of PrP fibrils co-exist and compete under homogenous conditions, which were hidden in bulk analyses of amyloid kinetics. The replication environment shapes the equilibrium between competing isomorphs. Different fibrillar isoforms form the basis of prion strains. Our data suggest that competing strains; indeed prions may exist in the form of quasispecies rather than a single isomorphic structure. Structural and dynamic polymorphism, may also underlie competition of infectious prions with non-infectious PrP amyloid *in vivo.* Single-particle analysis uniquely enables us to map the dynamic equilibrium between these species, which allows us to better understand the intricate balance of protein misfolding, amyloid formation and prion replication and which can inform more specific therapeutic interventions in the future.

## Acknowledgements

We thank Dr. Matthew Lew and Dr. Tianben Ding, Washington University in St. Louis, for help with computational analysis, Adam Wenborn and Dr. Jonathan Wadsworth, MRC Prion Unit, for providing mouse scrapie material and mass spectrometry, and Mark Batchelor, MRC Prion Unit, for help in protein expression.

## Author contributions

Conceptualization, J.B. and Y.S.; Methodology, J.B. and Y.S.; Investigation, Y.S., K.J., T.E., D.S.; Formal Analysis: Y.S.; Writing – Original Draft, Y.S., J.B.; Writing – Review & Editing, J.B., Y.S.; Funding Acquisition, J.C., J.B.; Resources, L.H., D.S.; Supervision, J.B, J.C.

## STAR Methods

### Resource availability

#### Lead contact

Further information and requests for resources and reagents should be directed to and will be fulfilled by the lead contact, Jan Bieschke.

#### Materials availability

This study did not generate new unique reagents or plasmids.

#### Data and code availability

All data and all original code is publicly available at Mendeley Data as of the date of publication. DOIs are listed in the key resources table. Any additional information required to reanalyse the data reported in this paper is available from the lead contact upon request.

### Method details

#### Generation of seeds for elongation assay

Frozen aliquots of monomeric PrP were thawed and diluted into buffer to a final composition of 10 μM rPrP in aggregation buffer (50 mM Na-phosphate pH 7.4, 2 M GdnHCl, 300 mM NaCl) with 20 μM ThT. The protein solution was pipetted into a 96-well plate (Corning 3651) with 3 Zirconium spheres inside each well. The plate was sealed and inserted into the plate reader (BMG Clariostar). Spontaneous aggregation was initiated by shaking the plate at 700 rpm with 100 s on / 20 s off cycles at 42 °C. ThT fluorescence was recorded every 10 min to monitor aggregation kinetics. The assay was stopped at 68 h when the kinetics of selected wells had reached plateau. Selected samples were collected.

Subsequently, a seeding assay was conducted, with the end product of the previous assay as seeds. Seed solution was prediluted 1:10, and sonicated for 10 min in a water bath sonicator. Then seed was added at 0.1% to protein solution whose composition was the same as the previous assay. Seeded aggregation was conducted under the same condition for spontaneous assay. Aggregates were collected after two days, aliquoted and used as seeds for elongation assays.

#### Single-particle elongation experiments

An 8-well microscope chamber was washed with Hellmanex II detergent and was plasma cleaned. Diluted seeds were incubated on the coverslip surface for 30 to 45s, allowing the seed particles to deposit onto the surface. After removing the excess seed, 200 μL monomeric PrP solution at a desired composition with 400 nM Nile Blue was added. The chamber was then sealed and ready to be imaged by the Nikon, Eclipse Ti2-E inverted microscope.

The temperature was maintained by an incubator box around the microscope body. For time -lapse imaging, Metamorph imaging software was used to automatically take images of multiple field of views (usually > 10) initially selected by users at a 15 min interval, for a total of > 2 days.

For data analysis, images taken at each location were imported into ImageJ as a time-lapsed image stack. The image stack was drift corrected by ImageJ plugin StackReg (Thevenaz et al., 1998) or Image Stabilizer (Li, 2008). Usually, 3-4 stacks were analyzed, and in total >70 fibrils that met our selection criteria were analysed for each set of conditions.

To extract kinetic information, a kymograph was generated for each fibril in ImageJ and the fibril edge was selected manually by drawing a segmented line. The saved fibril edge positions were analysed using custom scripts written in MATLAB, to identify the growth/stall phase, calculate the overall rate (final length/total time), pause free rate (final length/time spent in the growing phase), and stall percentage and other parameters.

#### Sequential seeding assay

A sequential seeding assay of MoPrP 91 was performed in buffer containing either 1 M (condition a) or 2 M GdnHCl (condition b), plus 50 mM Na-phosphate pH 7.4, 300 mM NaCl. Solution monomer concentration was 1 μM and seed concentration was 0.1% (w/w) with respect to monomer.

In the first seeding assay, the mixed solution with seed, monomer in the specific buffer was pipetted into plate wells with 3 Zr beads in each well, and incubated in BMG plate reader at 37 °C with agitation. The seeded products were labelled based on their buffer conditions, a or b, and were used as seeds for the second assay.

In the second assay, the use of two seed types (seed a and b) and two solution conditions (buffer a and b) generated four experiment combinations and four end products: aa, ab, ba and bb. Here the first letter represented the seed type (i.e. the buffer type of the first seeding assay), and the second letter represented the buffer condition of the second assay.

