## Supplementary Data and Methods for "Direct Observation of Competing Prion Protein Fibril Populations with Distinct Structures and Kinetics"

### Supplemental methods

#### *Protein expression and purification*

Recombinant truncated mouse prion protein PrP 91-231 (MoPrP 91) was expressed in Escherichia coli and purified using a protocol modified from (Jackson et al., 1999). Bacteria culture was grown at 37 °C with 100 µg/mL ampicillin and expression of His-tagged MoPrP 91 was induced by adding IPTG to 1 mM when the culture reached an OD600 of ~0.6. After growing overnight at 37 °C, cells were harvested by centrifugation. Cell pellet was resuspended in extraction buffer (50 mM Tris-HCl pH 8.0, 200 mM NaCl, 0.1% Tween-20, 50 U/ml benzonase, 10 µg/ml lysozyme) and disrupted by sonication. The suspension was centrifuged at 9900 g for 30 min and the resulting pellet was washed again in extraction buffer by resuspension, sonication and centrifugation. Solubilisation of protein was performed by resuspending the pellet in solubilisation buffer (6 M GdnHCl, 50 mM Tris-HCl pH 8.0, 0.8% β-mercaptoethanol), followed by disruption by sonication and centrifugation at 21,000 g for 45 min. The supernatant was saved and the pellet was solubilized again. Supernatants from both solubilisation steps were pooled and filtered through a 0.45-µm filter. The solution was loaded onto a 30 ml Ni-NTA column pre-equilibrated with buffer A (10 mM Tris, 100 mM $Na_2HPO_4$, 6 M GdnHCl, 10mM GSH pH 8.0). Oxidative refolding was carried out on column by running a 30 CV 0% - 100% buffer A to buffer B (10 mM Tris, 100 mM $Na_2HPO_4$, pH 8.0). After refolding, the protein was eluted with a linear gradient elution from buffer B to buffer C (10 mM Tris, 100 mM $Na_2HPO_4$, 1 M imidazole, pH 5.8) and protein fraction was collected. Imidazole was removed from the collected protein fraction by dialysis into 25 mM Tris pH 8.4. The His-tag was removed by Thrombin cleavage at room temperature overnight, in the presence of 2.5 mM $CaCl_2$. The cleaved protein was loaded onto a HiPrep SP FF cation exchange column pre-equilibrated with 10 mM HEPES pH 8.2. Protein was eluted with a linear gradient of 0-1 M NaCl. The eluted fraction was dialyzed against 10 mM HEPES buffer (pH 7.5), aliquoted and frozen for storage.

Monomeric PrP was prepared by spinning the protein through a 100 kDa membrane filter (Amicon). The flow-through was aliquoted and frozen.

*Purification of prion rods*

*Ex vivo* prions were purified from scrapie-infected mouse brains as previously described (Wenborn et al., 2015). Briefly, 10% (w/v) brain homogenate was pre-treated with Pronase, EDTA, sarcosine and Benzonase. NaPTA was added to 0.3%. After incubation, NaPTA precipitation was performed in the presence of 35% iodixanol. Pellets (containing contaminant proteins) and surface layer (comprised mainly of lipid and contaminant proteins) were discarded. The supernatant was filtered through a 0.45-µm microcentrifuge filter and diluted two-fold with 0.3% NaPTA+2% sarkosyl. After centrifugation, prion rods were enriched in the pellet. The pellet was washed twice by resuspending in 17.5% iodixanol+0.3% NaPTA and centrifugation. After the last centrifugation the pellet was resuspended in 0.1% sarcosine, aliquoted and frozen for storage. No PK digestion was performed.

Work with animals was performed under the licence granted by the UK Home Office (Project Licences 70/6454 and 70/7274) and conformed to University College London institutional and ARRIVE guidelines.

*Negative stain electron microscopy*

2 µL samples were loaded onto carbon-coated 300 mesh copper grids (AGS 160-3, AGAR Scientific) that were glow discharged for 40 seconds using PELCO easiGLOW™ glow discharge unit (Ted Pella Inc, USA). Samples were left to bind for 2 minutes, blotted, washed briefly in four water drops with blotting in between, and then stained with NanoW stain (Nanoprobes) 2 × 1 min, then blotted and air-dried. Grids were imaged on an FEI Talos electron microscope.

*Circular dichroism spectroscopy*

CD spectra were obtained using a Jasco J-715 spectropolarimeter (Jasco) in the wavelength range of 190-250 nm using a 2 mm Quartzglass Suprasil® cuvette (Hellma®, #110-2-40) with PTFE stopper lids. The scan rate was set at 100 nm/min, data pitch 0.5 nm with 2 nm band-width.

The temperature was set by a PTC-423L Peltier controller (Jasco Corporation, Japan). Prior to each measurement, the sample was left to equilibrate for 60 s. Ten scans were taken and averaged for each spectrum, which was background corrected by subtracting a buffer blank spectrum.

Chemical denaturation of $PrP^C$ was studied by titration experiments using a 6.4 M GdnHCl solution containing 50 mM NaP pH 7.4 and 300 mM NaCl at 37°C and ellipticity was monitored at 222 nm. The first measurement was acquired using the sample containing 10 µM $PrP^C$ in buffer with NaP and NaCl (no GdnHCl); a total of 23 measurements were obtained following the sequential addition of 25 µl of Guanidine until a final concentration of 4.4 M was reached. Once the denaturant was added, the solution was gently mixed four times and the sample was allowed to adjust for 60 seconds prior to every measurement. A total of three titrations experiments were performed in order to obtain reliable data.

The sample used for thermal denaturation was 5 µM $PrP^C$ in aggregation buffer. Thermal denaturation was studied between 10°C and 60°C by monitoring the ellipticity at 222 nm. The temperature interval was set at 0.1°C and the heating rate was 1°C/min.

*Preparation of microscope chamber*

8-well cell culture chambers with #1.5 high-performance glass coverslip bottom (Cellvis, 170 ± 5 µm thickness) were used for imaging. To remove fluorescent contaminants and to facilitate seed adhesion chambers were immersed in 2% Hellmanex II solution for 2-3 h at room temperature with occasional stirring, rinsed thoroughly with $H_2O$, followed by immersion in methanol for 5~10 min and rinsed with $H_2O$. The cleaned chambers were left on bench to dry, and then sealed with parafilm and stored in a closed container. Before each imaging or elongation experiment, chambers were glow discharged for 2 min using PELCO easiGLOW™ glow discharge unit.

*Chamber assembly for elongation experiments*

To prepare the monomer solution, monomeric PrP was diluted in the specific buffer to reach a desired final composition. Nile Blue was added to 400 nM for visualization of PrP fibrils.

Aliquoted recombinant seeds were thawed and diluted into aggregation buffer at a 1:80 or 1:120 dilution ratio. 10 µL diluted solution was deposited onto the coverslip surface inside one well of an 8-well microscope chamber and incubated for 30 to 45 s. The dilution ratio and incubation time were experimentally determined so that seed density on the coverslip surface was optimal. For authentic prion fibrils, the sample was incubated for 5-10 min. After incubation, the seed solution was removed and the well was washed twice with 200 µL aggregation buffer. 200 µL protein monomer solution in aggregation buffer was pipetted into the well. The chamber was sealed by double-wrapping in parafilm or by sealing the chamber lid with Twinsil silicone (Picodent). The sealed chamber with sample inside was then ready to be imaged under microscope.

*Optical Instrumentation*

Samples were imaged on a bespoke microscope system (Cairn Research Ltd., UK) based on a Nikon, Eclipse Ti2-E inverted microscope with a 100X objective (Nikon, CFI SR HP Apochromat TIRF 100XC/1.49 NA Oil) and iLas spinning TIRF illumination (Gattaca System). Lasers used for illumination in this work were 561 nm (OBIS LS laser) to visualize Nile Red fluorescence and 638 nm excitation laser (Omicron LuxX. Diode Laser) for Nile Blue fluorescence. A dual collimator was used allowing for quick switching between high intensity and large field imaging mode. TAB imaging utilized the high-intensity mode and time-lapse imaging of elongation utilized the large field imaging mode. Emission fluorescence passed through appropriate filters and was captured by a Photometric 95b CMOS camera. An enclosure box (OKO-Labs) built around the main body of the microscope maintained a constant temperature of the sample. MetaMorph software installed on the PC was used to control image acquisition.

*Time-lapse imaging of fibril elongation*

The assembled chamber with surface-attached seeds and monomer solution containing Nile Blue was placed on the microscope for ~20 min to equilibrate. Positions of multiple field of views (FOVs) with identified bright seeds were chosen using the multi-dimensional acquisition module in Metamorph imaging software. The software recorded images of those FOVs every 15 min, usually at 5% laser power at large field mode with 1 s exposure time.

After acquisition, images taken at each location were imported into ImageJ as a time-lapsed image stack. The image stack was drift corrected by ImageJ plugin StackReg (Thevenaz et al., 1998) or Image Stabilizer (Li, 2008). The middle 1000x1000 pixel-sized region was used for analysis. Usually, fibrils in 3-4 stacks were analyzed and the total fibril count was >70 for each set of conditions.

To eliminate bias in the selection of fibrils, all fibrils in a stack were analyzed, except for fibrils that met one of the following criteria: a) fibrils with contrast too low to distinguish them from the background; b) fibrils growing from a large seed cluster, i.e. more than 4 fibrils growing from one point into different directions; c) fibrils that intersected with other ones during growth; d) fibril ends that wobbled in the image plane or into solution; e) fibrils that went out of focus at later time points; or f) when an abrupt growth > 9 pixels (990 nm) took place between two adjacent time points, as it could not be excluded that these fibrils had grown in solution and were deposited between image acquisition.

To extract kinetic information, a kymograph was generated for each fibril by drawing a segmented line along its axis and running reslice command in ImageJ. The fibril edge in the kymograph was drawn as a segmented line and its coordinates were saved. The saved fibril edge positions were analysed using custom scripts written in MATLAB, to identify the growth/stall phase, and calculate the overall rate (final length/total time), pause free rate (final length/time spent in the growing phase), stall percentage and other parameters. All scripts are deposited in Mendeley Data http://dx.doi.org/10.17632/7mk7fxgzn4.1.

At least 70 fibrils were analysed for each experiment to generate a statistically robust dataset. The distributions of overall rates, pause-free rates, and stall percentages were plotted.

Fibril brightness was extracted from each kymograph. Since brightness fluctuated during fibril growth, the time point to take intensity measurement was set to be the end time of the last growth event to compare intensities between fibrils. If a fibril was in the growth phase when the experiment stopped, brightness was measured in the last frame. The brightness value was calculated as the average pixel intensity of the middle half of the fibril's length.

Since laser illumination was not flat but rather adopted a Gaussian profile in each FOV, measured fibril brightness needed to be corrected for the difference in illumination intensity, depending on the fibril's location in the FOV. To correct for the uneven illumination >200 points were selected from one FOV at background positions, i.e. at positions where no fibrils existed. Intensity values and (x,y) coordinates of selected points were fitted to a 2D Gaussian function. A brightness correction factor was calculated for any point (x,y) to be $G_{max}/G(x,y)$, where $G_{max}$ was the maximum value of the fitted Gaussian function, and G(x,y) was the value at coordinates (x,y). Thus, corrected fibril brightness was calculated by multiplying the brightness value by the correction factor at the fibril's (x,y) coordinates. Here we assumed that fibril intensity was linearly related to background intensity.

For one experiment where the illumination profile was not a regular Gaussian function (caused by a imperfect TIRF calibration), FOV was split into 10x10 grids. The correction factor of each grid was calculated from the mean value of selected background points in the grid and the maximum of those mean values in all grids.

*TAB imaging*

TAB imaging was performed for fibrils elongated on coverslip surfaces. Chamber assembly was the same as in 'Chamber assembly for elongation experiments' except that solution contained a reduced concentration of dyes to ensure better blinking; we used 50 nM Nile Red for imaging recombinant fibrils or 100 nM Nile Blue for imaging elongated authentic RML and ME7 fibrils. After incubation, elongated fibrils were imaged using 561 nm (for NR) or 638 nm (for NB) excitation laser at full laser power at high-intensity mode. Stacks of 5,000 or 10,000 frames with 20 ms exposure time were taken for each region of interest.

The captured image stacks were analyzed using ThunderSTORM plugin (Ovesný et al., 2014) in ImageJ to generate information on all blinking events. Reconstruction was further generated using a Matlab script with the brightness of each pixel representing the number of localizations that fell inside that pixel.

*Seeding on EM grid*

EM grids used for seeding assay were carbon-coated Au grids (EMS CF300-Au) as they were inert in the aggregation buffer. Grids were glow discharged for 40 s right before use. Recombinant seeds were diluted at 1:3.2 in buffer. 2 µL sample was pipetted onto the grid and incubated for 5~10 min in a closed container to prevent evaporation. The remaining sample was blotted. The grid was washed briefly in two buffer drops with blotting in between and put into a well of a 96-well plate (Corning 3651). 100 µL PrP monomer solution was added to the well. The plate was sealed and incubated in a 37 °C incubator for the indicated period of time.

After incubation, the solution was pipetted out from the well and the grid was taken out carefully, washed with three water droplets and stained with NanoW stain (Nanoprobes) 2 × 1 min, then blotted and air-dried. Grids were imaged on an FEI Talos electron microscope. Care was taken when handling grids as they were easily bent and carbon film peeled off easily during a multiple-step experiment.

Typically, fibrils longer than 500 nm were analyzed to obtain the width distribution. Fibrils imaged by TEM were straightened in ImageJ and laid horizontally. A 10-pixel wide, vertical line was drawn across each fibril, and an intensity profile along the line (average of intensities of 10 pixels covered by line width) was generated by ImageJ. Fibril edges were assumed to be at two dip positions and fibril width was calculated accordingly.

*CD sample preparation after sequential seeding assay in plate*

To prepare seeded samples for CD, samples aa and bb were used (1 ml each), and each was spun at 16,100 g for 2 h at RT. 700 µL supernatant was taken out and saved for CD measurements. Pellet was resuspended and sonicated for 15 min in a water-bath sonicator, and a further 10 min before CD measurements.

5 µM PrP monomer solutions in either of the two buffers used in samples aa and bb, respectively, were prepared as control samples. CD spectra were recorded for the buffer, monomer, pellet resuspension of aggregates and supernatant of aggregates. Spectra were then blank corrected by subtracting the corresponding buffer spectrum. Spectra for the two pellet resuspension samples were further baseline corrected. Finally, all spectra were smoothed using Adaptive-Smoothing filtering with a 5 nm window in Jasco software.

*Sequential surface seeding assay*

The first elongation assay in the microscope chamber was carried out with surface-attached seeds and 200 µL solution containing 1 µM PrP monomer, 1 M GdnHCl, 50 mM NaP pH 7.4, 300 mM NaCl, 400 nM NB. After incubating at 37 °C for 3 days, the chamber was opened, the solution taken out and the surface washed with buffer. Fresh monomer solution with 2 M GdnHCl (the other components were the same as in the previous assay) was added into the same well. The chamber was sealed again and put on objective for time-lapse imaging.

Intensity difference within a FOV caused by gaussian illumination needed to be adjusted. A pseudo background image was generated in ImageJ by running the Gaussian blur command with a radius of 50 pixels on the original image. Corrected image was obtained using Calculator plus command by 'divide' operation for each pixel: $i = (i_1/i_2)*k_1+k_2$, where $i_1$ was original image, $i_2$ was background image, $k_1$ = mean intensity of $i_2$, $k_2 = 0$.

*LCO assay*

Samples measured by LCO assay were $PrP^C$ in buffer with 1 M or 2 M GdnHCl, a, b, aa, ab, ba, bb. ThT-free samples were generated, as outlined above, and mixed with h-FTAA (LCO dyes, which were supplied by the Nilsson lab (Klingstedt et al., 2011)) to a final dye concentration of 1 µM. Samples were mixed thoroughly and dispensed into a 96-well plate in triplicate, with a minimum volume of 50 µL. The plate was left to incubate for ~30 minutes and then placed into a Tecan M1000 plate reader. 3D fluorescence maps (where both excitation and emission are varied) were recorded for relevant intervals. The data was exported into an excel spreadsheet and analysed in R studio using a script (adapted from (Lapworth and Kinniburgh, 2009)).

**Supplementary Figures**

**Figure S1.** MoPrP 91 characterization and seed generation. (A) MoPrP 91 aggregation kinetics measured by ThT fluorescence. The gray curve represents the kinetics of spontaneous aggregation. End-point aggregates were used as seeds at 0.1% concentration (w/w) for a sequential aggregation assay. The kinetics of seeding assays are shown by the orange and blue curves. End products were diluted and used as seeds for surface elongation assay. The two samples (represented by orange or blue curves) were generated in different experiments but essentially under the same conditions. Seeds represented by the blue kinetic trace were used for elongation assay studying the influence of $PrP^C$ concentrations; seeds represented by the orange traces were used for experiments studying GdnHCl series and temperature series. (B) typical TEM images of seeds used for elongation assays. Seeds were diluted before being absorbed onto EM grids; scale bar 200 nm. (C-F) MoPrP 91 denaturation curves. (C) Change in mean residue molar ellipticities during GdnHCl denaturation. (D) Fraction of unfolded $PrP^C$ monitored by CD spectroscopy and fitted curve. (E) $PrP^C$ thermal denaturation unfolding (blue dots) and refolding ellipticity curves (orange dots). (F) Fraction of unfolded $PrP^C$ during thermal unfolding.

**Figure S2.** Morphologies of PrP fibrils elongated in situ. (A) TAB images of fibrils elongated in 2 M GdnHCl. Scale bar 5 µm. (B) TEM images of fibrils elongated in 2 M GdnHCl. Scale bar 500 nm. (C) TEM images of fibrils elongated in 1 M GdnHCl. Scale bar 500 nm.

**Figure S3.** Kinetics of *in situ* elongation assay in PrP concentration series. (A) Distributions of fibril rate (left column), pause-free rate (middle column) and stall percentage (right column) under different $PrP^C$ concentration. (B) Stall percentages of three fibril types in monomer concentration series. Red line in each plot represents the average stall percentage. (C) Scatter plot of fibril brightness versus pause-free rate showing the grouping of type I (blue), II (orange) and III (purple) fibrils at each $PrP^C$

concentration. Ungrouped fibrils were shown as black circles. Figure relating to main manuscript Figure 5.

**Figure S4.** Kinetics of *in situ* elongation assay in GdnHCl concentration series (A) distribution of fibril rate (left column), pause-free rate (middle column) and stall percentage (right column) at different GdnHCl concentrations as indicated. (B). Scatter plot of fibril brightness versus pause-free rate showing the grouping of type I (blue), II (orange) and III (purple) fibrils. (C) pause-free rate distribution of type I, II and III fibrils at different GdnHCl concentrations, with fitted Gaussian curve (red curve). (D) histograms showing stall percentage of type I, II and III fibrils at different GdnHCl concentrations. Figure relating to main manuscript Figure 5.

**Figure S5.** Kinetics of *in situ* elongation assay in temperature series (A) distribution of fibril rate (left column), pause-free rate (middle column) and stall percentage (right column) at individual temperature. (B) Scatter plot of brightness versus pause-free rate showing the grouping of type I (blue), II (orange) and III (purple) fibrils. (C) pause-free rate distribution of type I, II and III fibrils at different temperatures with fitted Gaussian curve (red curve). (D) histograms showing stall percentage of type I, II and III fibrils at each temperature with mean value (red line). Figure relating to main manuscript Figure 5.

**Figure S6.** Determination of repeating patterns in single fibril intensity profiles from TAB images of PrP fibrils seeded by RML (A) or ME7 (B) prion rods, respectively. Fibrils were traced in ImageJ; intensity plot profiles and average distances between intensity peaks ± SD are shown for each fibril.

Supplementary Figure 1

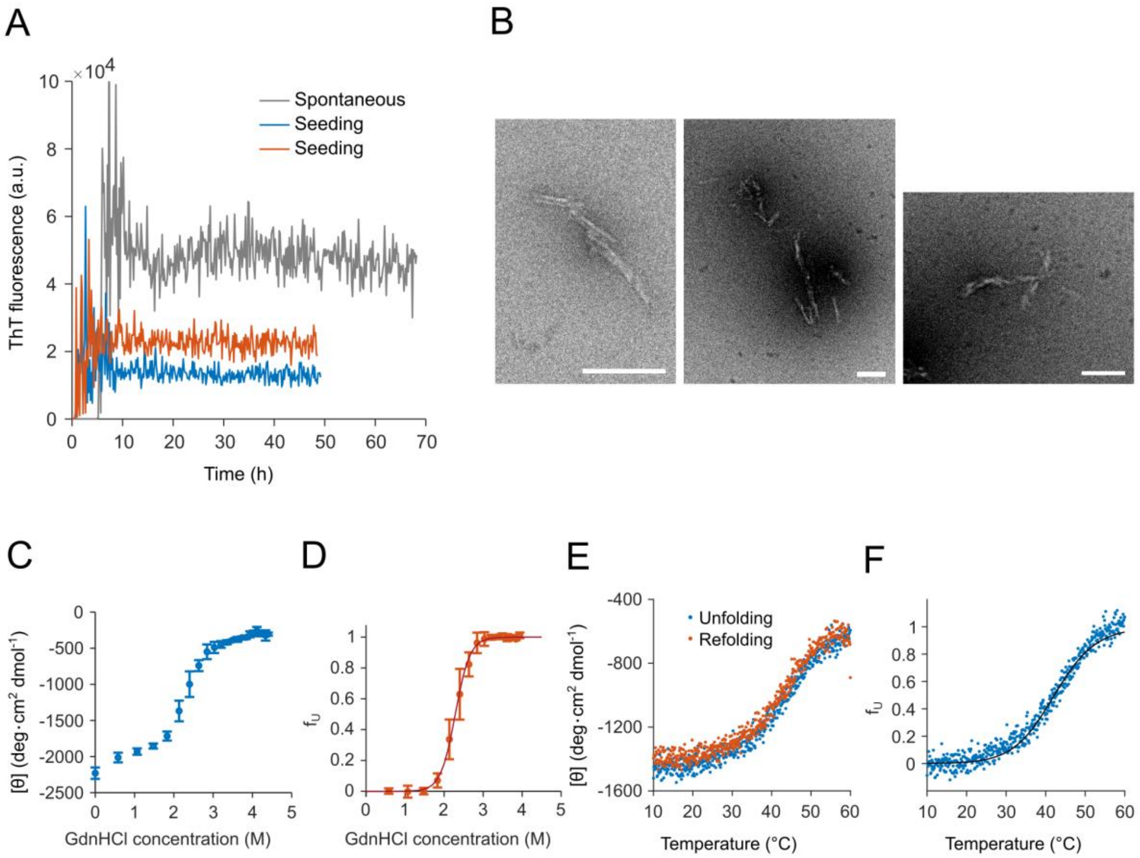

Supplementary Figure 2

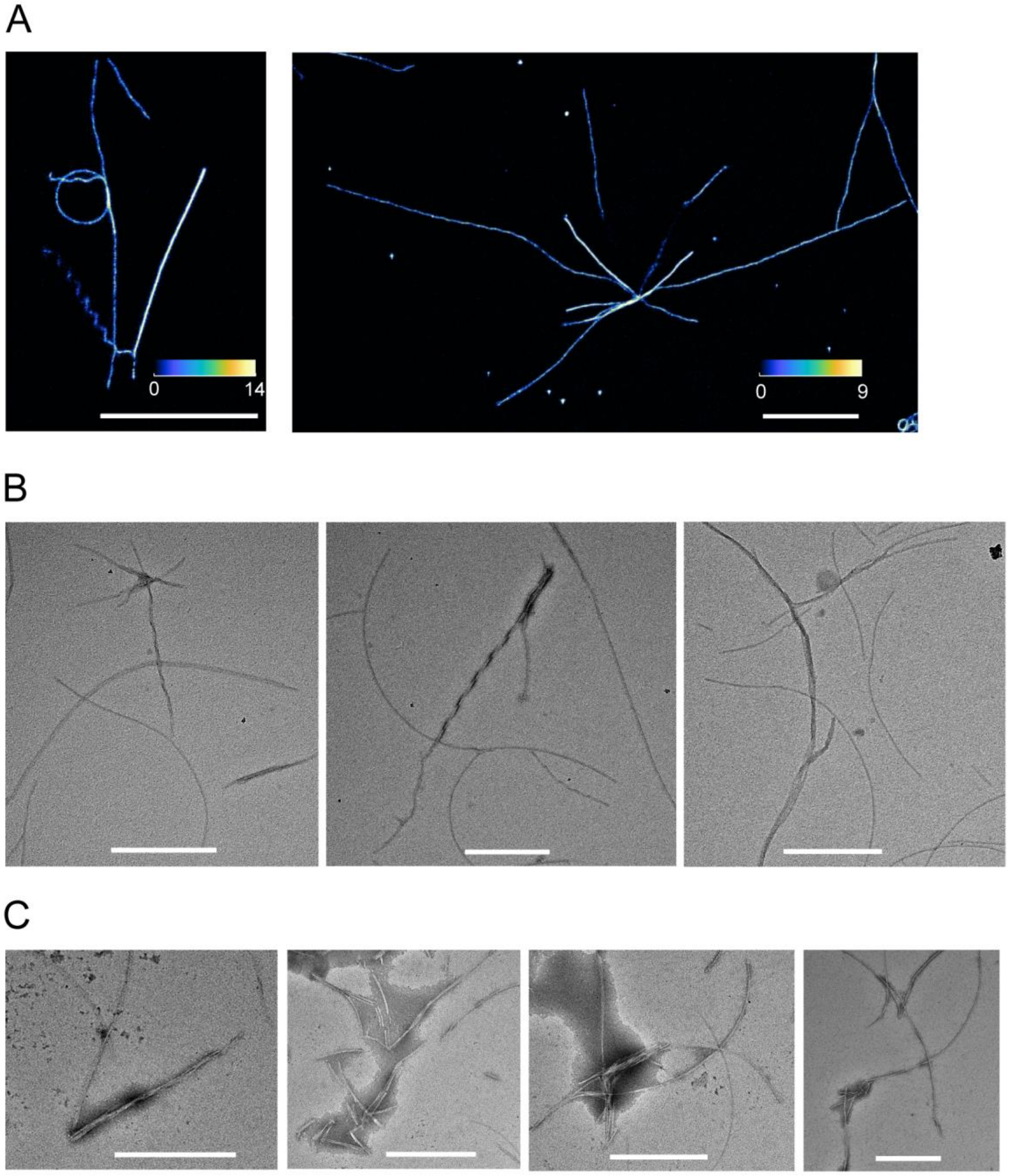

Supplementary Figure 3

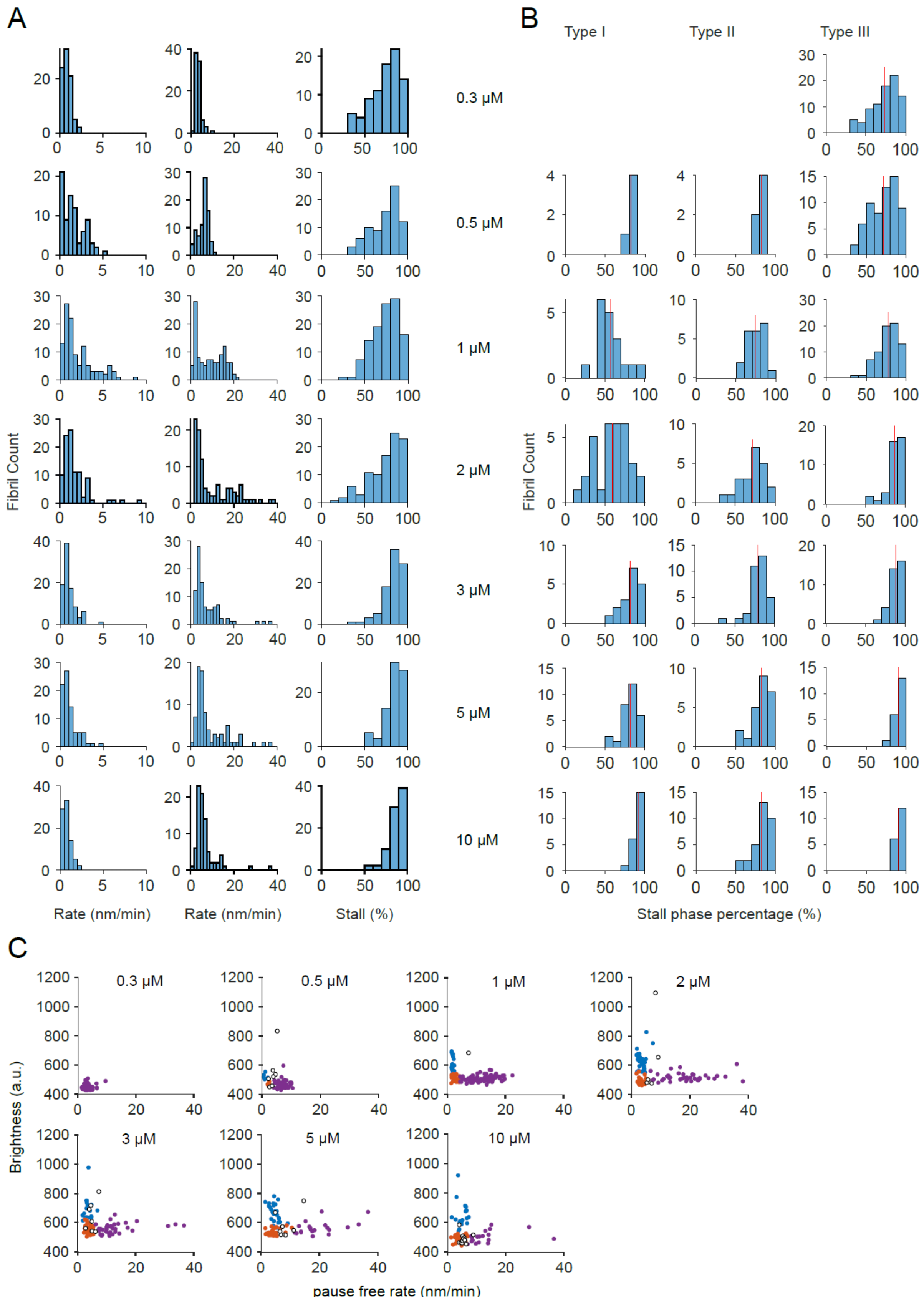

Supplementary Figure 4

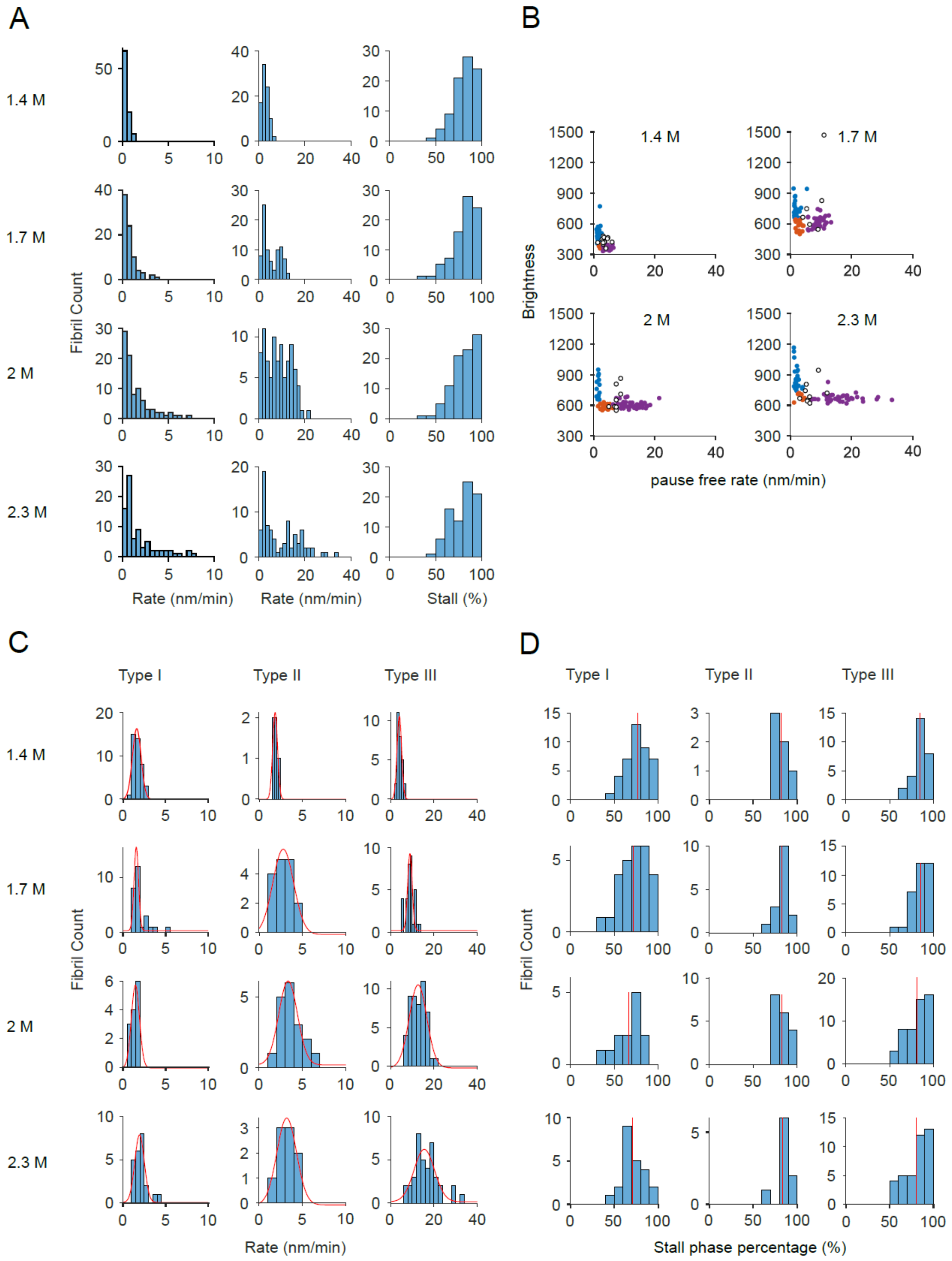

Supplementary Figure 5

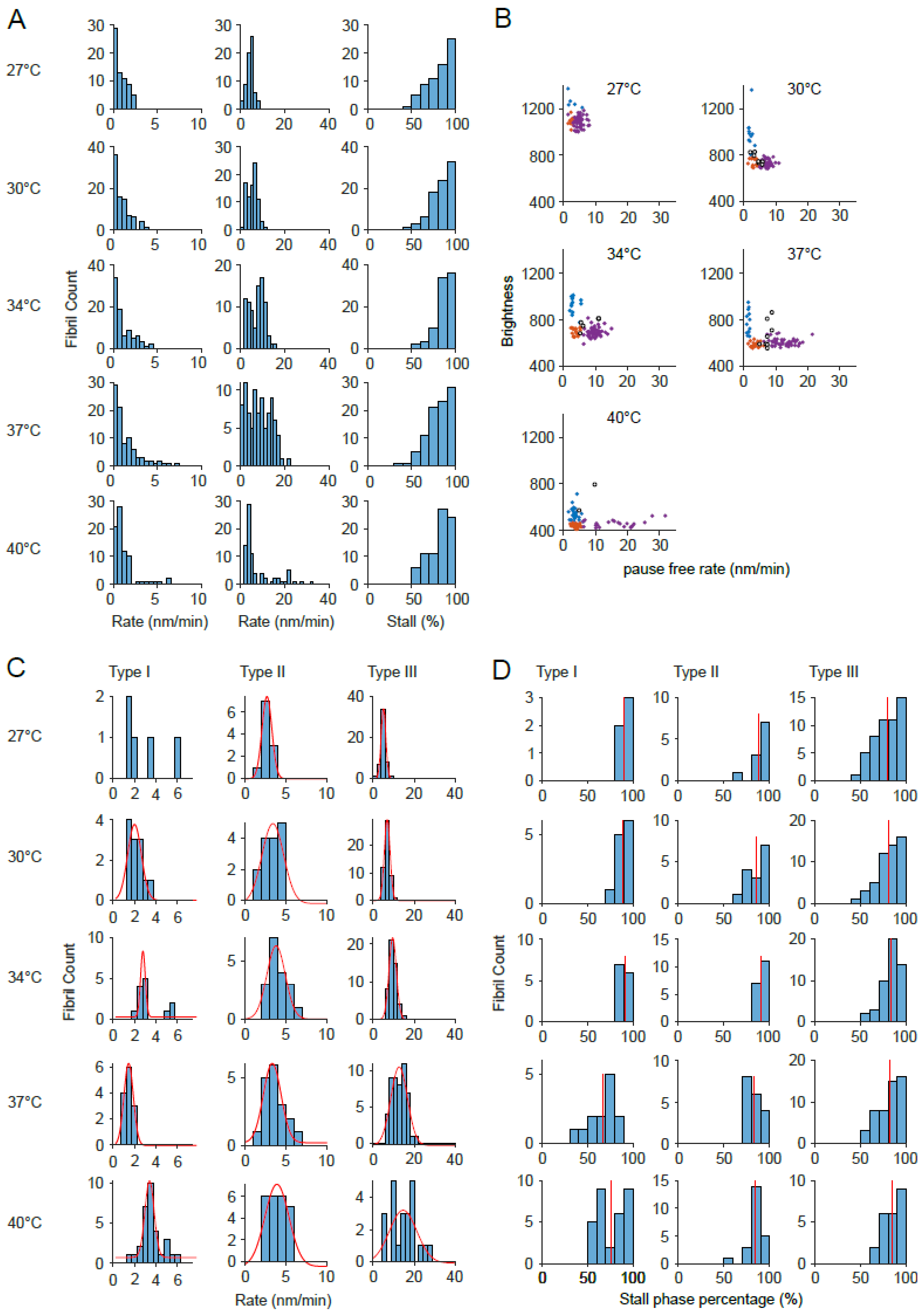

Supplementary Figure 6

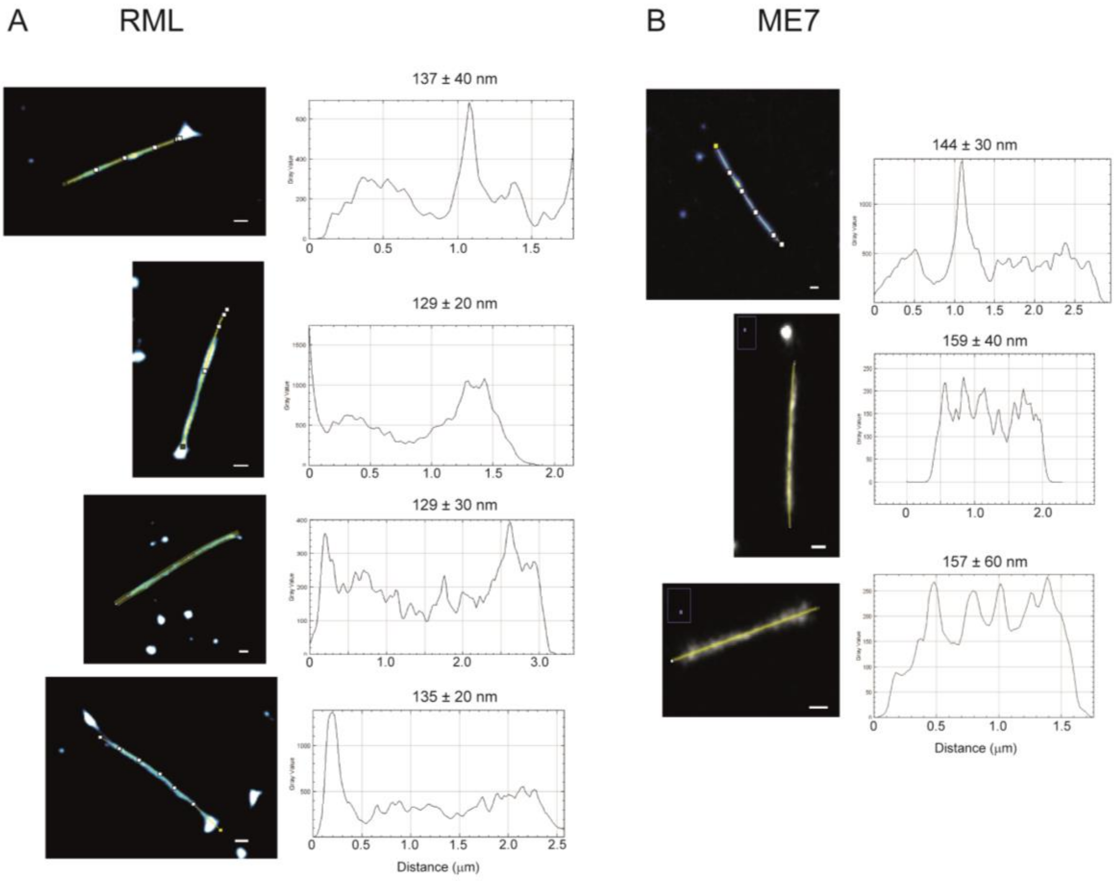
